# Small nuclear RNAs enhance protein-free RNA-programmable base conversion on mammalian coding transcripts

**DOI:** 10.1101/2024.06.12.598766

**Authors:** Aaron A. Smargon, Deepak Pant, Sofia Glynne, Trent A. Gomberg, Gene W. Yeo

## Abstract

Endogenous U small nuclear RNAs (U snRNAs) form RNA-protein complexes responsible for eukaryotic processing of pre-mRNA into mature mRNA. Previous studies have demonstrated the utility of guide-programmable U snRNAs in targeted exon inclusion and exclusion. We investigated whether snRNAs can also enhance conversion of RNA bases over state-of-the-art RNA targeting technologies in human cells. When compared to adenosine deaminase acting on RNA (ADAR)-recruiting circular RNAs, we find that guided A>I snRNAs consistently increase adenosine-to-inosine editing efficiency for genes with higher exon counts, perturb substantially fewer genes in the transcriptome, and localize more persistently to the nucleus where ADAR is expressed. A>I snRNAs can also edit pre-mRNA 3′ splice sites to promote splicing changes. Finally, snRNA fusions to H/ACA box snoRNAs (U>Ψ snRNAs) increase targeted RNA pseudouridylation efficiency. Altogether, our results advance the protein-free RNA base conversion toolbox and enhance minimally invasive RNA targeting technologies to treat genetic diseases.

## INTRODUCTION

Recently the gene editing field has turned from CRISPR (Clustered Regularly Interspaced Short Palindromic Repeats) and other exogenous protein-encoded multicomponent systems toward minimally invasive single-component guided RNA scaffolds that recruit highly expressed endogenous protein machinery to edit genes at the RNA level. Researchers have particularly focused on suppressing in-frame premature termination codons (PTCs) caused by single base-pair substitution nonsense mutations in coding regions of mRNA transcripts. PTCs, which account for an estimated 10-15% of human genetic diseases such as Cystic Fibrosis and Hurler syndrome, lead to truncated proteins and subsequent degradation of PTC-harboring mRNAs by nonsense mediated mRNA decay^1^.

Given a clearly defined mechanism, several minimally invasive strategies already exist to treat PTC-associated diseases. More clinically established drugs like splice-switching antisense oligonucleotides and small molecules could be administered to patients^2–4^, but not all PTC diseases are amenable to exon skipping and PTC-suppressor small molecules lack target site specificity. Similarly, engineered suppressor tRNAs designed to read through PTCs at the translational level could do so at any targeted stop codon context sequence^5, 6^. In contrast to these approaches, programmable guided RNA scaffolds that recruit highly expressed endogenous proteins to convert PTC bases directly strike a safer balance between minimal invasiveness and specificity.

One such class of systems recruits endogenous adenosine deaminase acting on RNA (ADAR) enzymes to convert PTC adenosines to inosines (A-to-I). In mammalian cells, active ADAR family members ADAR1 and ADAR2 recognize regions of nuclear double-stranded RNA (dsRNA), primarily at Alu repetitive regions but also in coding regions and even at splice sites^7^. After A-to-I conversion, splicing and translation machinery generally recognizes inosine as its structurally comparable base, guanosine (G). Leveraging this finding, several groups have encoded a cytosine (C)-mismatch guided RNA scaffold of both linear and circular form which, when hybridized to the RNA sequence surrounding a targeted adenosine, recruits endogenous ADARs that efficiently convert the targeted adenosine opposite the cytosine to inosine^8–10^. While robust editing is possible, ADARs display a strong preference for the UAG motif, with diminished activity for the other PTC sequence contexts of UGA and UAA, in addition to varying by cell type expression^11^.

Another class of systems utilizes H/ACA box snoRNPs, ribonucleoproteins highly conserved across eukaryotes which catalyze the uridine-to-pseudouridine (U>Ψ) conversion on snRNAs, ribosomal RNAs (rRNAs), and some mRNAs^12, 13^. In mammalian cells, H/ACA snoRNAs recruit four core proteins, DKC1, NOP10, NHP2, and GAR1. Together, these proteins convert U-to-Ψ at a site between two guide-templated regions specified by the H/ACA snoRNA. Building upon initial work performed in yeast^14^, two groups reprogrammed human H/ACA snoRNAs to convert U-to-Ψ at all three PTC sequence contexts (UAG, UGA, and UAA), leading to successful translational readthrough^15, 16^. While a promising approach, snoRNA localization predominantly to the nucleolus, and not to the nucleoplasm where pre-mRNAs are transcribed and processed into mRNAs, has limited its base-conversion potential.

Although the gene editing field has evolved beyond CRISPR, important lessons endure from CRISPR’s success. For example, a peptide nuclear localization signal (NLS) was critical for translation of Cas9 activity from prokaryotic to eukaryotic cells^17, 18^. Analogously, we hypothesized that achieving optimal subcellular localization of programmable guided RNA scaffolds could enhance the ability of endogenous protein machinery to perform base conversion on target coding RNA transcripts. To test this hypothesis, we selected as a putative RNA nucleoplasm localization signal components of endogenous Uridine-rich small nuclear RNAs (U snRNAs), which natively recruit splicing machinery to process pre-mRNA into mature mRNA and have previously been engineered to modulate RNA splicing^19, 20^. In this new application, we evaluated the capacity of U snRNAs to enhance protein-free base conversion, both A-to-I and U- to-Ψ, on mammalian coding transcripts.

## RESULTS

Preclinical studies utilizing engineered U1 and U7smOPT snRNAs have already shown promise for the inclusion and exclusion of exons to treat disease^19, 20^. In fact, an AAV9-mediated U7smOPT snRNA gene therapy to treat boys with DMD exon 2 duplications is currently in Phase I/II clinical trials (ClinicalTrials.gov ID: NCT04240314). Based on this established track record, and the fact that most other U snRNAs are recruited downstream of U1, we concentrated on these two U snRNAs. U1 snRNAs (bound by highly expressed U1A, U170K, U1C, and members of the Sm core) initiate the major spliceosome to splice introns out of pre-mRNA. Meanwhile, U7 snRNAs initiate the 3′ end processing of non-polyadenylated histone pre-mRNAs. Researchers previously mutated U7 snRNAs into U7smOPT snRNAs that bind only the Sm core and not LSm proteins. With backbone sizes of 153nt and 45nt respectively, U1 and U7smOPT snRNAs are comparatively small and easily encodable in a variety of genetic delivery vehicles, from lipid nanoparticles to adeno-associated virus.

### A-to-I editing with engineered U snRNAs

Of the existing single-component A>I programmable guided RNA scaffolds, circularized ADAR-recruiting RNAs (cadRNAs) have demonstrated potent editing with a simple design^9, 10^. Due to their elegant circularization by autoligating twister ribozymes, cadRNAs effectively withstand degradation by exoribonucleases to sustain good expression in cells. cadRNAs contain a C-mismatch guide with typically 100nt homology regions flanking either side of the mismatched C and occasionally mismatches and loops throughout these flanking regions to inhibit spurious bystander editing by ADAR. Given that spliceosomal component Sm proteins have been found to associate with ADAR1 and ADAR2, programmable U snRNAs may possess inherent A-to-I editing capacity^21^. To test this conjecture, we replaced the cadRNA backbone (in a U6 promoter/U6 terminator cassette) with either the U7smOPT backbone (in a U7 promoter/U7 terminator cassette) or U1 snRNA backbone (in a U1 promoter/U1 terminator cassette) at the 3′ end of fixed C-mismatch guides (Fig. 1a).

**Figure 1:**
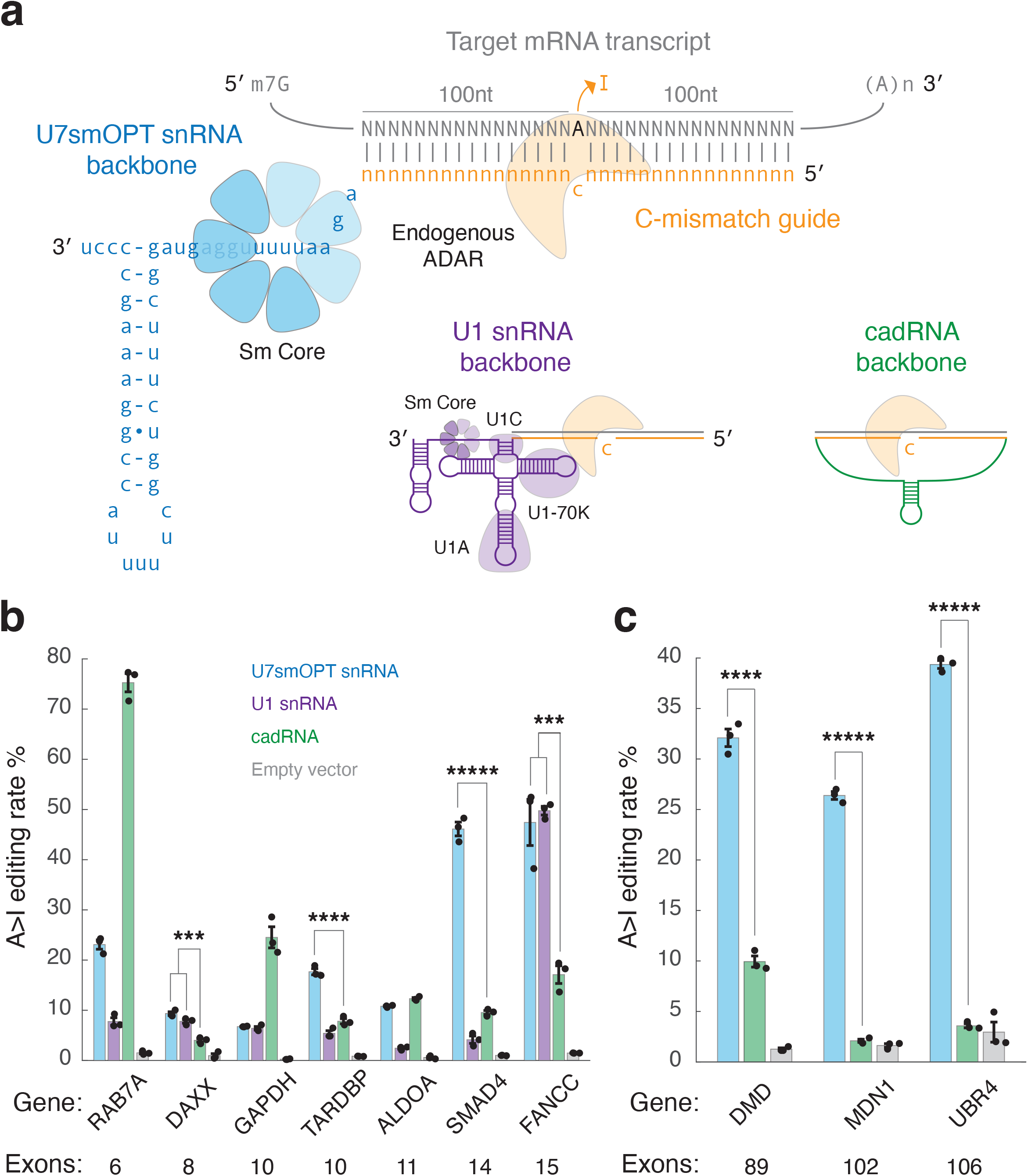
Programmable U snRNAs convert A-to-I on endogenous human transcripts. **a**, Schematic of three different RNA-guided A>I base converters targeting an mRNA transcript with the same C-mismatch guide: U7smOPT (blue), U1 snRNA (purple), and cadRNA (green). **b,** Editing percent performance by transfection in HEK293T cells of three A>I base converters on targets sites from various human genes. A>I overperformance significance vs. cadRNA: **\*\*\***, **\*\*\*\***, **\*\*\*\*\***: *p* < 1e-3, 1e-4, 1e-5 (one-way ANOVA, Bonferroni correction for multiple comparisons). Error bars reflect standard error of mean. **c,** Editing percent performance by transfection in HEK293T cells of U7smOPT snRNA and cadRNA A>I base converters on targets sites from human genes with high exon counts. A>I overperformance significance vs. cadRNA: **\*\*\*\***, **\*\*\*\*\***: *p* < 1e-4, 1e-5 (one-way ANOVA). Error bars reflect standard error of mean.

A head-to-head A-to-I editing test of the two U snRNAs against cadRNA across seven endogenous loci in HEK293T cells revealed a few interesting findings (Fig. 1b). First, U1 snRNA almost invariably performed more poorly than U7smOPT snRNA. We reasoned that its greater molecular complexity and proclivity for splicing machinery recruitment caused this limiting effect, and so we disregarded U1 snRNA for the remainder of our study. Second, although U7smOPT snRNA bested cadRNA across four of the seven loci, neither appeared a clear winner. Finally, U7smOPT snRNA most unambiguously outshined cadRNA editing performance at SMAD4 and FANCC, loci for genes with the highest exon counts. Taking this notion to its logical extreme, we tested the backbones on three loci of human genes with among the highest exon counts: DMD, MDN1, and UBR4 (89, 102, and 106 exon counts, respectively) (Fig. 1c). Here U7smOPT snRNA convincingly outperformed cadRNA across all high exon count gene loci. Given that high exon count genes tend to be larger and more prone to accumulating disease-relevant mutations (as is the case for DMD where ∼15% of Duchenne muscular dystrophy-implicated mutations are nonsense)^22^, U7smOPT snRNAs present an attractive new modality for treating PTC diseases.

### Off-target genetic perturbations of A>I snRNAs

Next, we asked how U7smOPT snRNAs compared to cadRNAs with respect to off-target genetic perturbations. Selecting one guide where cadRNA outperformed (RAB7A targeting) and one where U7smOPT snRNA outperformed (DMD targeting), we performed differential gene expression analysis with DESeq2 on two replicates of RNA sequencing data from each condition compared to empty vector (significance cutoffs of |log2 (fold change)| > 0.5 and adjusted *p*-value < 0.05) (Fig. 2a, Supplementary Fig. 1)^23^. In analyzing the data, we removed apparent overexpression of DMD due to a library preparation artifact as has been done in previous work (Supplementary Fig. 2)^9^. In either case, U7smOPT snRNA produced far fewer genetic perturbations (∼4-fold to 8-fold) than did cadRNA, both in genes upregulated and downregulated. Pathway analysis with Metascape of perturbed genes conserved across both cadRNA conditions and absent from either U7smOPT snRNA condition showed notable downregulation of Herpes simplex virus 1 infection and double-strand break repair via synthesis strand annealing (Supplementary Fig. 3)^24^. These results suggest that cadRNAs may be inducing an innate immune response and acting as templates for homologous recombination, either of which would be highly problematic for cells.

**Figure 2:**
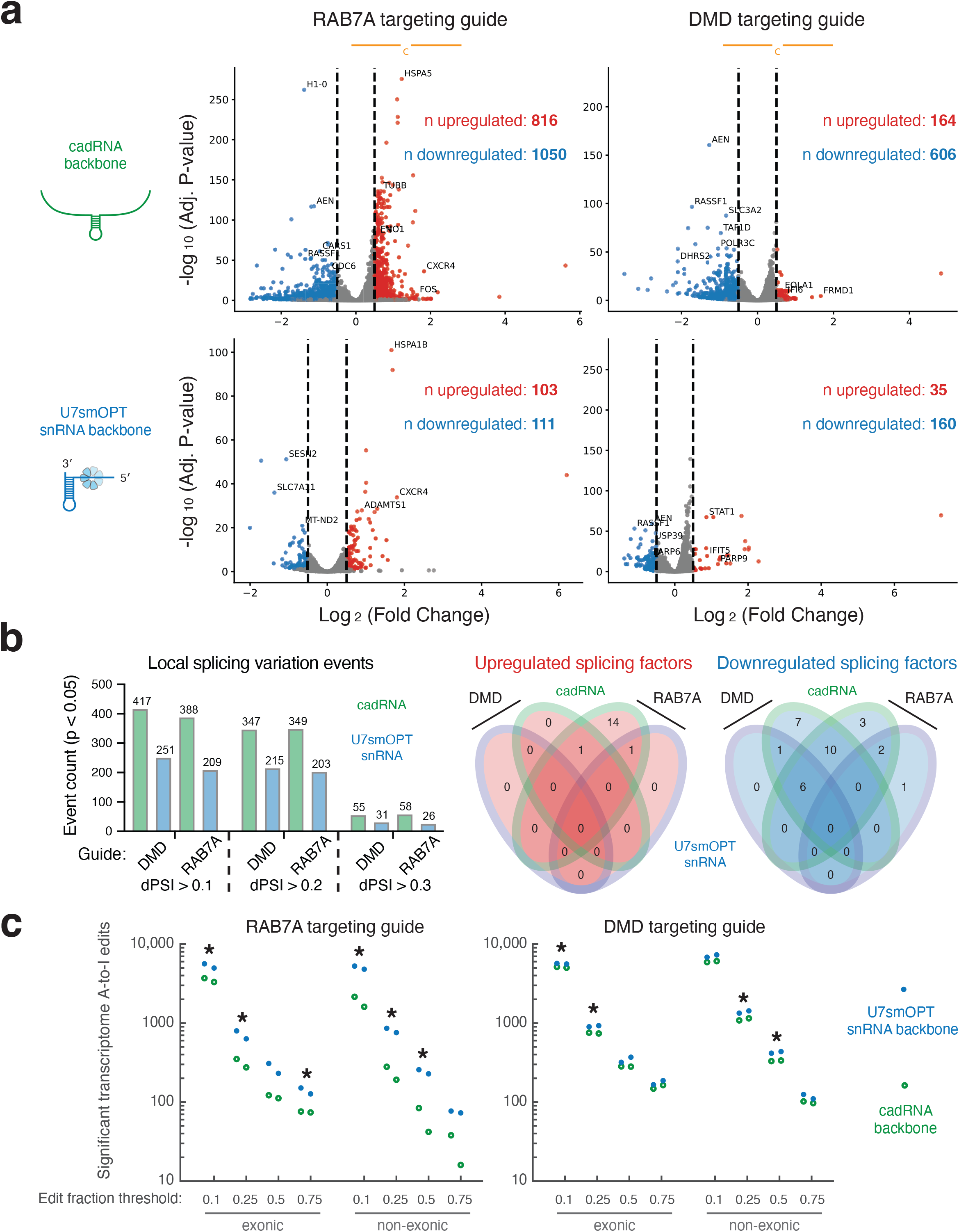
A>I snRNAs perturb fewer genes than circularized ADAR-recruiting RNAs. **a**, Scatterplots of differential gene expression analysis against empty control (pUC19) of cadRNA backbone vs. U7smOPT snRNA backbone for RAB7A- and DMD-targeting guides. Cutoffs for significance are |log2 (fold change)| > 0.5 and adjusted *p*-value < 0.05. Upregulated genes are colored in red, and downregulated genes are colored in blue. **b,** Number of local splicing variation events from differential splicing analysis against empty control (pUC19) of cadRNA backbone vs. U7smOPT snRNA backbone for RAB7A- and DMD-targeting guides, with *p*-value < 0.05 and different thresholds of differential Percent Spliced In (dPSI) for events (left). 4-way Venn diagrams of the number of significantly upregulated and downregulated splicing factors represented in the significantly perturbed genes from **a (**right). **c,** Counts of significant transcriptome A-to-I edits, both exonic and non-exonic, absent in both empty control (pUC19) condition replicates of cadRNA backbone vs. U7smOPT snRNA backbone for RAB7A- and DMD-targeting guides, with different edit fraction thresholds (1 = 100% editing). Edit count significance of U7smOPT vs. cadRNA: *p* < 5e-2 (one-way ANOVA).

While cadRNA may be more genetically perturbative overall, we expected U7smOPT snRNA to generate more splicing changes in the transcriptome. To test this hypothesis, we performed local splicing variation (LSV) analysis with MAJIQ of the RNA sequencing data sets (significance cutoff of *p*-value < 0.05) (Fig. 2b, Supplementary Fig. 4)^25^. Astonishingly, for both guides across three different thresholds of differential Percent Spliced In (dPSI), cadRNA produced ∼1.5x to ∼2x more significant LSV events than did U7smOPT snRNA. We attribute this unexpected finding not to directly guided splicing perturbations, but rather to pleiotropic effects stemming from cadRNA-mediated downregulation of splicing factors (24 vs. 7 for DMD targeting and 21 vs. 3 for RAB7A targeting, cadRNA vs. U7smOPT snRNA, respectively).

Finally, we examined the number of off-target A-to-I editing events absent from both empty vector replicates and present across cadRNA and U7smOPT replicates with our established SAILOR pipeline (significance cutoff of >75% confidence) (Fig. 2c). For both guides—for both exonic and non-exonic edit sites, and at various edit fraction thresholds—U7smOPT snRNA generated consistently more transcriptome-wide A-to-I edits than did cadRNA.

### Pre-mRNA base editing with A>I snRNAs

We reasoned that the seeming contradiction between higher cadRNA-mediated genetic perturbations and U7smOPT-snRNA transcriptome-wide A-to-I edits could be reconciled by known more durable expression of cadRNAs (and accompanying antisense knockdown of transcripts) coupled with higher localization of U7smOPT snRNAs to the nucleoplasm where ADAR proteins are expressed. For not only do U snRNAs spend most of their life cycle in the nucleoplasm^26^, but also circular RNAs are actively exported to the cytosol^27^. This localization hypothesis would also explain why U7smOPT outperforms cadRNA on high exon count gene mRNAs, which typically persist longer in the nucleus due to more extensive splicing prior to nuclear export.

To test the localization hypothesis, we devised a subcellular localization quantitative polymerase chain reaction (qPCR) assay whereby qPCR performed on both A>I scaffolded guides and genes from nuclear and cytosolic fractionated RNA enables inferred nuclear:cytosolic ratio comparison between U7smOPT snRNA and cadRNA for equivalent guides (Fig. 3a). As expected, across three different guides U7smOPT localized more highly to the nucleus than did cadRNA, with a sample-matched NEAT1 positive control showing no significant difference in nuclear:cytosolic ratio between conditions (Fig. 3a).

**Figure 3:**
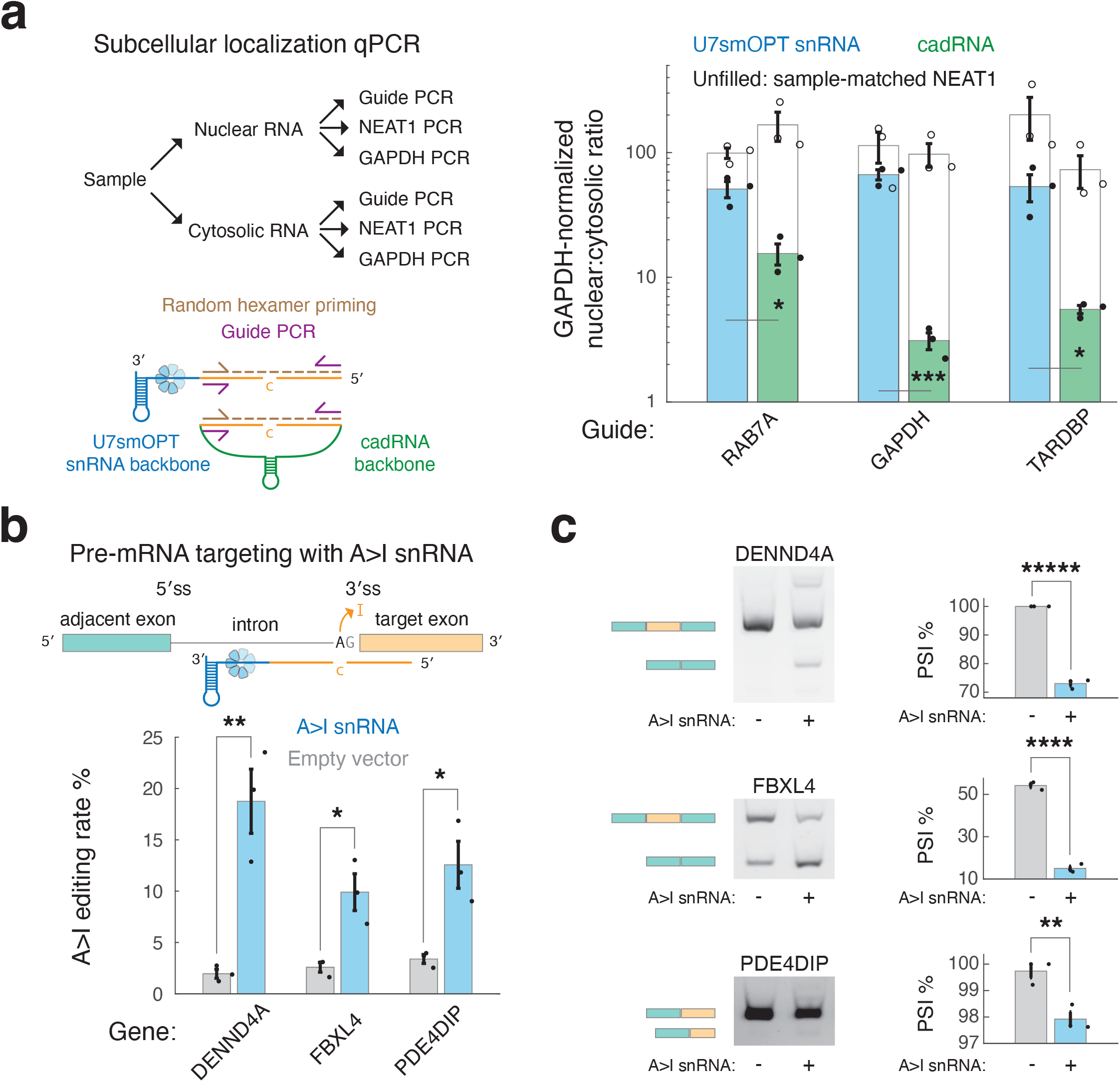
Targeted nuclear editing of pre-mRNA splice sites by A>I snRNAs alters mRNA splicing. **a**, Schematic of subcellular localization qPCR to determine nuclear:cytosolic enrichment of U7smOPT snRNA and cadRNA A>I base converters (left). Results from subcellular localization qPCR of the A>I base converters with guides targeting three different genes, and the lncRNA NEAT1 as a positive control for the assay (right). Nuclear:cytosolic enrichment significance vs. cadRNA: **\***, **\*\*\***: *p* < 5e-2, 1e-3 (one-way ANOVA). Error bars reflect standard error of mean. **b,** Schematic of A>I snRNA targeting 3′ splice site of pre-mRNA (top). Editing percent performance by transfection in HEK293T cells of A>I snRNA on 3′ splice sites of three human genes (bottom). A>I editing significance vs. empty vector: **\***, **\*\***: *p* < 5e-2, 1e-2 (one-way ANOVA). Error bars reflect standard error of mean. **c,** RT-PCR gels with quantified Percent Spliced In (PSI) of experiments from **b**. PSI significance vs. empty vector: **\*\***, **\*\*\*\***, **\*\*\*\*\***: *p* < 5e-2, 1e-4, 1e-5 (one-way ANOVA). Error bars reflect standard error of mean.

Given this exciting discovery, we wondered whether A>I snRNAs (C-mismatch guides with U7smOPT snRNA backbones) could be leveraged to edit pre-mRNAs, an application not robustly demonstrated with single-component A>I programmable guided RNA scaffold modalities. We tested A>I snRNAs on the 3′ splice sites (3′ss) of three pre-mRNAs for which native A>I editing has previously been implicated in splicing perturbation through ADAR knockdown: DENND4A, FBXL4, and PDE4DIP (Fig. 3b)^28^. More efficient U1 promoter-driven A>I snRNAs with 100nt homology regions flanking a mismatched C successfully edited all three pre-mRNA loci, with editing rates ranging from ∼10-20%. Importantly, all three A>I snRNAs resulted in splicing perturbation mirroring the kind (exon skipping vs. exon boundary change) and degree (tens of dPSI to ones of dPSI) of the previous study’s loci (Fig. 3c, Supplementary Figs. 5,6)^28^. Our results indicate a new use of A>I snRNAs whose efficacy may depend largely on cis-splicing factors.

### Increased pseudouridylation efficiency with U-to-Ψ snRNAs

Given our success in localizing A>I snRNAs to the nucleus for enhanced A-to-I editing, we applied a similar approach to U-to-Ψ RNA base conversion. H/ACA snoRNAs, which catalyze U-to-Ψ modification of rRNAs and snRNAs, localize predominantly to the nucleolus. We hypothesized that more nucleoplasmic localization of programmable guided H/ACA snoRNAs via fusion to a U7smOPT snRNA backbone would direct the snoRNAs away from the nucleolus for more efficient base conversion U-to-Ψ on coding RNAs (Fig. 4a).

**Figure 4:**
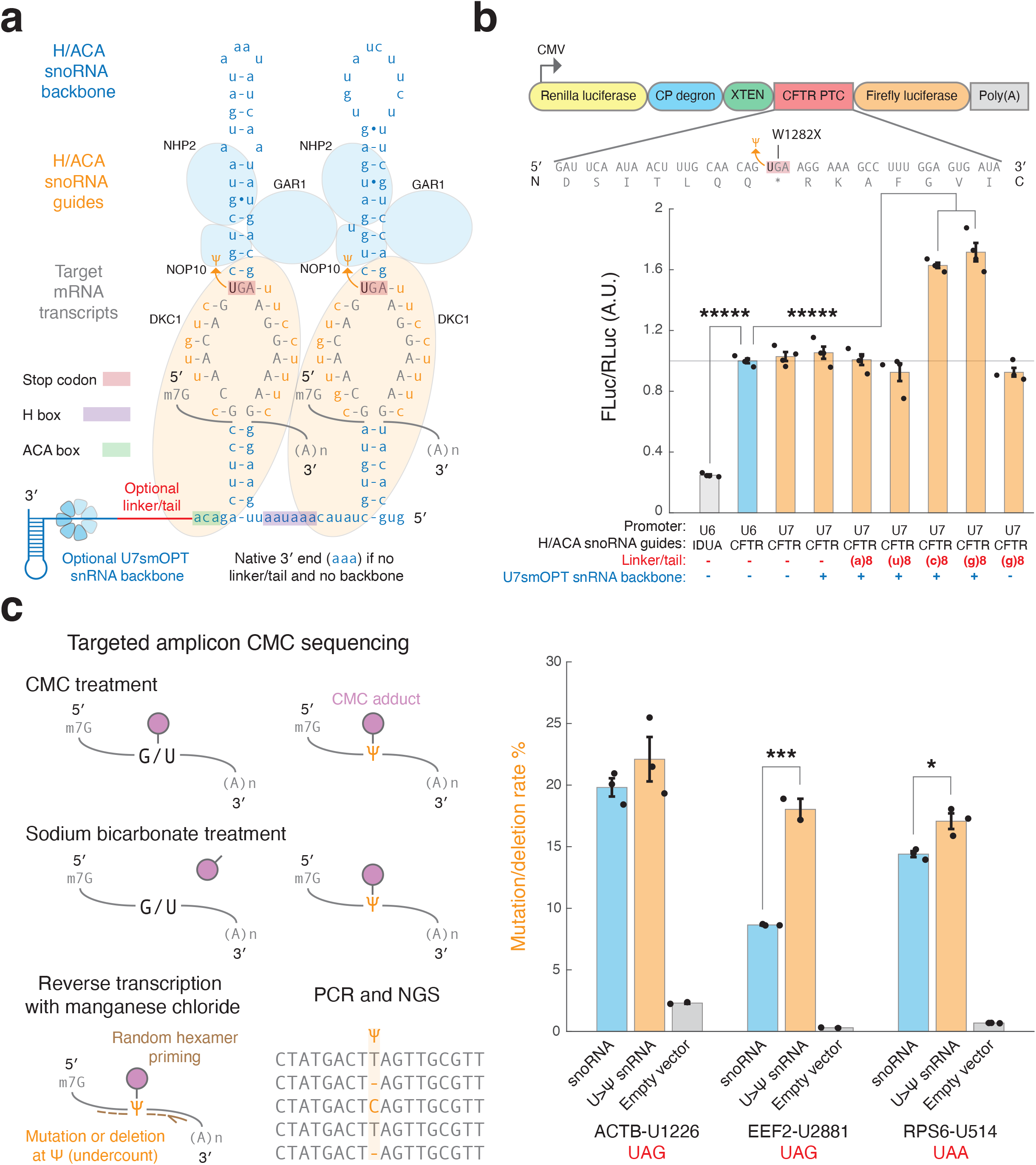
U>Ψ snRNAs demonstrate increased potency over engineered snoRNAs. **a**, Design of engineered U>Ψ snRNA, with guided H/ACA snoRNA targeting an mRNA transcript fused by RNA linker to U7smOPT backbone. **b,** Schematic of dual-luciferase reporter to quantify premature termination codon (PTC) suppression of CFTR mutant W1282X by tested U>Ψ snRNA designs (top). Luciferase ratio performance by co-transfection in HEK293T cells of various U>Ψ snRNA designs on PTC suppression dual-luciferase reporter (bottom). FLuc/RLuc significance vs. U6 promoter-driven CFTR guide H/ACA snoRNA without linker/tail and U7smOPT backbone: **\*\*\*\*\***: *p* < 1e-5 (one-way ANOVA, Bonferroni correction for multiple comparisons). Error bars reflect standard error of mean. **c,** Schematic of targeted amplicon CMC sequencing to infer pseudouridylation of endogenous targeted mRNA by mutation/deletion rate (left). Mutation/deletion rate performance by transfection in HEK293T cells of guided U>Ψ snRNAs vs. H/ACA snoRNAs on stop codon context sequences from three human genes (right). Mutation/deletion rate overperformance significance vs. snoRNA: **\***, **\*\*\***: *p* < 5e-2, 1e-3 (one-way ANOVA). Error bars reflect standard error of mean.

To test this hypothesis, we designed a monocistronic, internally controlled dual-luciferase reporter harboring a disease-implicated PTC from human CFTR (W1282X) between Renilla and Firefly luciferase (Fig. 4b). In a co-transfection experiment in HEK293T cells, expression of an established CFTR target-guided H/ACA snoRNA increased FLuc/RLuc luminescence ratio ∼4x over a negative control IDUA target-guided H/ACA snoRNA, validating the assay’s sensitivity as a proxy for PTC readthrough. Of all linkers tested between CFTR target-guided H/ACA snoRNA and U7smOPT snRNA backbone, both the (c)8 and (g)8 linkers (5′-cccccccc-3′ and 5′-gggggggg-3′) resulted in significant FLuc/RLuc luminescence ratio increases, with (g)8 linker raising the luminesce ratio by ∼70% but a (g)8 tail without U7smOPT snRNA backbone having no significant effect. In light of these results, we suspect that unlike the other linkers tested the (c)8 and (g)8 linkers help stabilize the expanded RNA scaffold by preventing RNA processing downstream of the ACA box.

Finally, we tested the H/ACA box snoRNA-(g)8 linker-U7smOPT snRNA backbone fusions (U>Ψ snRNAs) on three endogenous loci in HEK293T cells using an established targeted amplicon CMC (N-cyclohexyl-N′-β-(4-methylmorpholinium) ethylcarbodiimide) sequencing assay whose output mutation/deletion rate is generally regarded as an undercount of U-to-Ψ modification (Fig. 4c)^15^. On all three loci, two with statistical significance and one by over 100%, U>Ψ snRNAs outperformed H/ACA box snoRNAs as predicted.

## DISCUSSION

RNA base conversion by programmable single-component guided RNA scaffolds has demonstrated promise as both a minimally invasive and target-specific approach to gene editing. In this study, we engineered U snRNAs to enhance such systems for A-to-I and U-to-Ψ conversion on mammalian coding transcripts. In either base conversion case, snRNAs improved system safety and/or efficacy performance over the state-of-the-art with an aspiration toward preclinical targeted suppression of premature termination codon diseases. Given that Sm core proteins are highly conserved and expressed in all mammalian cells, we expect our findings to translate effectively to other cell types and broadly to PTC disease therapeutics.

More efficient genetically encodable single-component RNA-guided editing of RNA noncoding regions, including the 3′ splice sites of pre-mRNA, opens other therapeutic opportunities. For example, RNA base conversion snRNAs could edit intronic RNA-binding protein (RBP) motifs to displace destabilizing RBPs and increase nuclear RNA expression. When coupled with single-component RNA-guided translational activation systems, RNA base conversion snRNAs could provide an additional boost to protein expression^29^.

Importantly, snRNA enhancements to RNA base conversion systems are guide-independent, suggesting an approach that will benefit researchers even as they orthogonally optimize RNA guides for on-target base conversion efficiency and specificity. While our study focused on the U7smOPT snRNA backbone, there are likely further optimizations to be made to the sequence of the snRNA enhancement for augmented scaffold stability and nucleoplasmic RNA localization. In addition to the H/ACA snoRNA, we anticipate that snRNA components could enhance programmable base conversion by other noncoding RNAs, such as C/D box snoRNAs^30–33^. Future studies will undoubtedly explore these open questions and potential use cases.

## ONLINE METHODS

### Cloning of plasmids

Plasmids with guides/snoRNAs (Supplementary Tables 1,2) were subcloned into pUC19 (N3041S, NEB) and pcDNA 3.1(-) (V79520, Life Technologies Corporation), both digested with EcoRI-HF (R3101T, NEB) and BamHI-HF (R3136T, NEB), by Gibson Assembly Master Mix (E2611L, NEB) with specified sequences of oligonucleotides. Gibson assemblies were transformed in Mix & Go! Competent Cells-JM109 (T3005, Zymo Research Corporation) and plated on LB agar plates with antibiotic. Colonies were cultured in LB media with antibiotic, then miniprepped with QIAprep Spin Miniprep Kit (27106, Qiagen). Resulting miniprepped plasmid DNA was sequenced by Primordium Labs and verified by SnapGene.

### Cell culture

Human HEK 293T cells (Takara Bio, 632180) were maintained in D10 (DMEM (4.5 g/L D-glucose) supplemented with 10% Fetal Bovine Serum (Gibco) and 1% Penicillin-Streptomycin (10,000 U/mL) (Gibco) at 37 °C with 5% CO2. Cells were periodically passaged once at 70-90% confluency by dissociating with TrypLE Express Enzyme (Gibco) at a ratio of 1:10.

### Transfections

Plasmids were transfected into human HEK 293T cells under passage 30 by jetOPTIMUS DNA transfection Reagent (76299-632, VWR International). For A-to-I RNA extractions, cells were plated in 48-well tissue culture plates and transfected at ∼60% confluency with 250 ng of plasmid DNA. For U-to-Ψ RNA extractions, cells were plated in 12-well tissue culture plates and transfected at ∼60% confluency with 1 ug of plasmid DNA. For luciferase reporter assay, cells were plated in 96-well tissue c ture plates and transfected at ∼60% confluency with 100 ng of plasmid DNA (75ng of guide plasmid DNA, 25 ng of reporter plasmid DNA). Under these conditions, a transfection efficiency of 80+% was achieved routinely.

### RNA extraction, A>I editing quantification, and RNA sequencing library preparation

At a time 48 hours after transfection, cells were washed twice with PBS, then RNA was extracted by RNeasy Plus Mini Kit (74136, Qiagen) with an elution volume of 30 uL. For A>I editing quantification, cDNA synthesis was carried out by ProtoScript II First Strand cDNA Synthesis Kit (E6560L, NEB) with 3 uL RNA in a 10 uL total volume using oligo(dT) primers supplied with the kit. PCR with 500nM specified primers (Supplementary Table 3) was carried out by NEBNext Ultra II Q5 Master Mix (M0544L, NEB) with Tm of 68°C and 30 second extension time for 30 cycles for mRNA products and for 32 cycles for pre-mRNA products. PCR products were purified by QIAquick PCR Purification Kit (28106, Qiagen) and submitted to Sanger sequencing to quantify A>I editing. RNA sequencing library preparation was carried out by Illumina Stranded mRNA Prep, Ligation (20040532, Illumina) with 1 ug RNA using manufacturer’s standard protocol. The RNA sequencing library was sequenced on an Illumina NovaSeq X Plus 10B under the PE100 configuration with a target sequencing depth of ∼50 million reads per sample.

### RNA sequencing alignment

Reads were checked for quality and adapter sequences using FastQC. Paired reads were then aligned to the genome using STAR aligner version 2.7.6a. A genome index for alignment was generated using the GENCODE v.44 hg38 primary assembly with the following command line parameters: --genomeFastaFiles GRCh38.primary_assembly.genome.fa, --sjdbGTFfile gencode.v44.primary_assembly.annotation.gtf, and --sjdbOverhang 100. Paired FASTQ files were then mapped to the genome with default options and --outSAMtype BAM unsorted. Aligned reads in BAM format were sorted by position using samtools sort (version 1.3.1) with default parameters. Sorted BAM files were indexed using samtools index with default parameters.

### Differential gene expression analysis

Differential expression of aligned RNASeq reads relative to controls was analyzed using subreadfeaturecounts version 1.5.3 followed by DESeq2 version 1.39.3. Gene counts for control (pUC19) and experimental replicates were collected in count matrices using subreadfeaturecounts (v 1.5.3). Gene count matrices for each condition were generated using the function subreadfeaturecounts::featureCounts with the following parameters:

-p (for paired-end reads), -a gencode.v44.primary_assembly.annotation.gtf, -t exon, -g gene_name, --primary (to count only the primary alignment for multimapping reads) -Q 255 (the minimum quality score that a counted read must satisfy), and --ignoreDup (to exclude duplicate reads). Matrices were then loaded into a jupyter R notebook as a ‘counts’ object using R::read.table, excluding the first 5 lines which do not contain counts. Counts in the R data frames were labelled by condition followed by labels.df <-DataFrame(condition = labels, row.names = colnames(counts)). Count matrices were converted into DESeq2 datasets with DESeq2:: DESeqDataSetFromMatrix. Genes with no expression across samples were removed. Differentially expressed genes were then identified using DESeq2::DeSeq with default parameters. Results were collected and saved as .csv files using DeSeq2:: results and R:: write.table.

Volcano plots of significantly up and downregulated genes for each condition were generated using RNAlysis 2 (v 3.9.2). Genes from DESeq2 (v 1.39.3) outputs were plotted using DESeqFilter.volcano_plot() with alpha (p-adj significance threshold) set to 0.05 and Log2FC thresholds of 0.5 and 1.00.

An additional counts matrix containing gene counts for all experimental and control replicates was used to generate PCA plots with RNAlysis 2 (v 3.9.2). Counts were normalized by relative log expression with RNAlysis2::CountFilter.normalize_rle(). Genes with no expression across columns were filtered from analysis using CountFilter.filter_low_reads() with a threshold of 0. PCA was performed using RNAlysis2::CountFilter.pca() function with default parameters, except for power_transform = False.

GSEA (v 4.3.2) was used for pathway analysis of the ranked and filtered gene lists. DESeq2 output tables were copied into Microsoft Excel (v 16.66.1). Ranking metrics for each gene were calculated by the formula =SUM(SIGN([@log2FoldChange])*(-LOG10([@pvalue]))). Genes were then filtered by a LogFC threshold of +/- 0.5. Ranked lists containing gene names and ranking metrics were copied into .rnk files. For DMD targeting guide RNAs, DMD was excluded from lists due to a library preparation artifact. Ranked lists were loaded into GSEA and analyzed using GSEAPreRanked with the following parameters changed from default: gene set = c5.go.bp.v2023.2.Hs.symbols.gmt, collapse = no_collapse, Enrichment statistic = classic, and Create SVG plot images = true.

### Gene pathway analysis

Significantly upregulated and downregulated genes identified in both cadRNA datasets but not in either U7smOPT snRNA dataset were analyzed for gene pathway signficance with Metascape (v3.5.20240101) using as background all genes without an “NA” p-adj significance in all replicates of cadRNA and U7smOPT snRNA datasets from the DESeq2 pipeline.

### Differential splicing analysis

MAJIQ and VOILA software packages (v 2.5) were used to assess splicing variation between treatment groups (cadRNA and U7 smOPT) and pUC19 controls. MAJIQ builder constructed splice graphs, and MAJIQ quantifier was used to quantify delta percent spliced in (dPSI) of local splicing variations (LSVs) under default conditions at known splice sites. VOILA TSV function was used select only genes with at least one LSV with 95% > probability the dPSI > x ( P(|dPSI| > x) > 0.95). Plotting LSVs was performed with PRISM and R.

### Splicing factor analysis

Significantly perturbed splicing factors were identified as a subset of significantly upregulated and downregulated genes from both cadRNA and U7 smOPT datasets contained within GOBP_RNA_SPLICING (GO:0008380) at the Gene Ontology Consortium

### Transcriptome-wide A>I editing analysis

Transcriptome wide RNA editing was quantified with SAILOR (v 1.1.0). Base quality MD tags for aligned reads were generated using samtools (v 1.3.1) calmd with -b (for bam files) and the GENCODE v. 44 GRCh38.primary_assembly.genome.fa sequence file. Reads containing MD tags were then analyzed with the SAILOR (v 1.1.0) cwl workflow. The reference genome used was the same used for alignment and the addition of MD tags. Edit sites were only considered significant if their SAILOR confidence level was greater than 0.75. Edit sites were identified as exonic if they intersected by bedtools intersect command (parameters “-s -wa -a”) with features labeled as “exon” from gencode.v44.primary_assembly.annotation.gtf. All other edit sites were identified as non-exonic. Edit fraction thresholds (1 = 100%) were applied on SAILOR output POST_PSEUDOCOUNT_EDIT%.

### Subcellular localization qPCR

Cells were washed twice with ice cold PBS, then spun down for 5 minutes at 300 g, with supernatant aspirated. Cells were then resuspended completely by gentle pipetting with 150 uL Buffer A (15mM Tris-HCl pH 8, 15mM NaCl, 60mM KCl, 1mM EDTA pH 8, 0.5mM EGTA pH 8, 0.5mM spermidine, 10U/mL SUPERase•In RNase Inhibitor (AM2694, ThermoFisher Scientific)). To this solution was added and mixed by inversion 150 uL of 2x lysis buffer (Buffer A with 0.5% NP-40). Mixture was incubated for 8 minutes at 4°C, then spun down for 5 minutes at 400 g. The top 200 uL of supernatant was carefully removed and placed into a new tube (this is the cytosolic fraction). The remaining supernatant was removed and discarded from the nuclear pellet, and this pellet was resuspended in 1mL of RLN buffer (50mM Tris-HCl pH 8, 140mM NaCl, 1.5mM MgCl2, 0.5% NP-40, 10mM EDTA pH 8, 10U/mL SUPERase•In RNase Inhibitor (AM2694, ThermoFisher Scientific)). This nuclear resuspension was incubated for 5 minutes at 4°C. During the incubation, the cytosolic fraction was spun again for 1 minute at 500 g, and its supernatant was collected into a new tube. 500 uL Trizol LS (10296010, Invitrogen) was added to this cytosolic fraction. The nuclear fraction was spun down once more for 5 minutes at 500 g. Supernatant was removed from the nuclear fraction pellet, and 500 uL TRIzol (15596018, Invitrogen) was added to the nuclear fraction pellet. To both TRIzol homogenizations was added 1 uL of GlycoBlue Coprecipitant (AM9516, ThermoFisher Scientific). Then RNA was extracted by phenol-chloroform extraction, followed by ethanol precipitation.

cDNA synthesis was carried out by ProtoScript II First Strand cDNA Synthesis Kit (E6560L, NEB) with 6 uL RNA in a 20 uL total volume using random hexamer primers supplied with the kit. Prior to qPCR, nuclear fraction cDNA was diluted 1:2 with nuclease-free water, and cytosolic fraction cDNA was diluted 1:60 with nuclease-free water. qPCR with 500nM specified primers (Supplementary Table 3) was carried out by PowerTrack SYBR Green Master Mix (A46109, ThermoFisher Scientific) using the CFX Opus 384 (Bio-Rad) and qPCR parameters of 95°C for 2 minutes, followed by 40 cycles of 95°C for 15 seconds and 60C°C for 1 minute.

From the qPCR Cq values, subcellular (nuclear and cytosolic) guide and NEAT1 expression levels were normalized relative to their subcellular GAPDH expression controls, and then the ratio of these normalized subcellular expressions was calculated. Data were analyzed and plotted in MATLAB (v R2024a).

### Splicing isoform quantification

For splicing isoform quantification, cDNA synthesis was carried out by ProtoScript II First Strand cDNA Synthesis Kit (E6560L, NEB) with 3 uL RNA in a 10 uL total volume using oligo(dT) primers supplied with the kit. PCR with 500nM specified primers (Supplementary Table 3) was carried out by NEBNext Ultra II Q5 Master Mix (M0544L, NEB) with Tm of 68°C and 30 second extension time for 32 cycles. PCR products were purified by QIAquick PCR Purification Kit (28106, Qiagen), with 50% of eluted volume run on a E-Gel EX Agarose Gels, 2% (G402022, ThermoFisher Scientific) for ∼15 minutes with E-Gel Ultra Low Range DNA Ladder (10488096, ThermoFisher Scientific) and E-Gel 50 bp DNA Ladder (10488099, ThermoFisher Scientific). Gels were visualized using the Azure Biosystems c600. Gel bands were quantitated using GelAnalyzer (v19.1), with Percent Spliced In (PSI) values calculated after adjusting bands for relative molecular weights.

### Luciferase reporter assay

At a time 48 hours after transfection, cell media was changed, and luminescence was generated using the Dual-Glo Luciferase Assay System (E2920, Promega) according to manufacturer’s standard protocol. Luminescence was measured using a Tekan infinite 200Pro plate reader with the Costar 96 flat white setting, automatic attenuation, and integration times of 500ms for Firefly and 100ms for Renilla. Data were analyzed and plotted in MATLAB (v R2024a).

### Targeted amplicon CMC sequencing

To disrupt the RNA secondary structure, 5 ug of RNA per sample in ∼10 uL nuclease-free water was incubated at 80°C for 5 minutes, and quickly chilled on ice. Next, the denatured RNA was transferred into 100 uL of freshly prepared and sterile-filtered BEU buffer (50 mM Bicine, pH 8.5, 4 mM EDTA, 7 M urea) with 0.2 M CMC (C106402-1G, Sigma Aldrich) and incubated at 37°C for 20 min, followed by purification with ethanol precipitation. The RNA pellets were then dissolved in 50 uL Na_2_CO_3_ buffer (50 mM Na_2_CO_3_ pH 10.4, 2 mM EDTA) and incubated at 37°C for 2 hours, followed by ethanol precipitation. Next, the RNA pellets were dissolved in 10 nuclease-free water.

2 uL of random hexamer primers from SuperScript First-Strand Synthesis System for RT-PCR (11904018, ThermoFisher Scientific) were added, and the mixtures were denatured at 65**°**C for 5 min followed by chilling on ice. Next, 8 uL freshly prepared 2.5x reverse transcription buffer (125 mM Tris pH 8.0, 15 mM MnCl2, 187.5 mM KCl, 1.25 mM dNTPs, 25 mM DTT) was added to the RNA-primer mixtures and the mixture was incubated at 25**°**C for 2 min. Then 1 uL SuperScript II reverse transcriptase from SuperScript First-Strand Synthesis System for RT-PCR (11904018, ThermoFisher Scientific) was added, and the reactions were carried out at 25**°**C for 10 minutes, 42**°**C for 3 hours and 70**°**C for 15 minutes.

For library preparation, PCR was carried out in two steps. In the first step, PCR with 500nM specified locus-specific primers (Supplementary Table 3) was carried out with 1 uL of cDNA by NEBNext Ultra II Q5 Master Mix (M0544L, NEB) in a total reaction volume of 10 uL with Tm of 65°C and 30 second extension time for 15 cycles. Then first round PCR products were cleaned up by 1.8x AMPure XP beads (A63881, Beckman Coulter) and eluted in 11 uL. In the second step, PCR with 500nM specified NGS universal and barcoded primers (Supplementary Table 3) was carried out with 10 uL of cleaned-up first round PCR product by NEBNext Ultra II Q5 Master Mix (M0544L, NEB) in a total reaction volume of 20 uL with Tm of 65°C and 30 second extension time for 15 cycles. Second round PCR products were pooled and purified by QIAquick PCR Purification Kit (28106, Qiagen), then this purified pool was run on a E-Gel EX Agarose Gels, 2% (G402022, ThermoFisher Scientific) for ∼15 minutes. The upper band on the gel (∼250bp) was extracted and purified by QIAquick Gel Extraction Kit (28704, Qiagen) before sequencing. The targeted amplicon sequencing library was sequenced on an Illumina NovaSeq X Plus 10B under the PE150 configuration with a target sequencing depth of ∼5 million reads per sample.

### Targeted amplicon CMC sequencing analysis

Targeted amplicon sequencing FASTQ files were collapsed along UMI (Unique Molecular Identifier) by selecting as sequence for a given 10-mer UMI in a read its first occurrence in each FASTQ file. Then only sequences with a perfect 8-mer matches both upstream and downstream of target NΨ at the locus were considered. Deletion rates were calculated as the fraction of total sequences at a locus with 1 nucleotide between the perfect 8-mer matches. Mutation rates were calculated as the fraction of total sequences at a locus with 2 nucleotides between the perfect 8-mer matches and which did not have a T at the Ψ position in target NΨ.

## ACKNOWLEDGMENTS

We thank members and alumni of the Yeo laboratory, in particular Dr. Orel Mizrahi, Dr. Stefan Aigner, and Samuel Hatch for their input on the research, and Steven Blue for his support in lab. This publication includes data generated at the UC San Diego IGM Genomics Center utilizing an Illumina X Plus that was purchased with funding from a National Institutes of Health SIG grant (#S10 OD026929). G.W.Y. is supported by NIH R01 HG004659, U24 HG009889 and an Allen Distinguished Investigator Award, a Paul G. Allen Frontiers Group advised grant of the Paul G. Allen Foundation. A.A.S. was supported by a Biomedical Research Fellowship from the Hartwell Foundation.

## AUTHOR CONTRIBUTIONS

A.A.S. was primarily responsible for designing and executing experiments, analyzing data, and writing the manuscript, under the supervision of G.W.Y. D.P. assisted in experiments and analyses related to targeted amplicon CMC sequencing. S.G. and T.A.G. analyzed A>I RNA sequencing data. All authors interpreted data and reviewed the paper prior to publication.

## COMPETING INTERESTS

G.W.Y. and A.A.S. have filed for a patent related to this work. G.W.Y. is a co-founder, member of the board of directors, scientific advisory board member, equity holder and paid consultant for Locanabio and Eclipse BioInnovations. G.W.Y. is a visiting professor at the National University of Singapore. G.W.Y.’s interests have been reviewed and approved by the University of California, San Diego in accordance with its conflict-of-interest policies. The authors declare no other competing financial interests.

## DATA AVAILABILITY

RNA-seq data from this study will be made available during peer review at the National Center for Biotechnology Information’s Gene Expression Omnibus.

## REFERENCES

1. Mort, M., Ivanov, D., Cooper, D.N. & Chuzhanova, N.A. A meta-analysis of nonsense mutations causing human genetic disease. Hum Mutat 29, 1037–1047 (2008).

2. Havens, M.A. & Hastings, M.L. Splice-switching antisense oligonucleotides as therapeutic drugs. Nucleic Acids Res 44, 6549–6563 (2016).

3. Howard, M., Frizzell, R.A. & Bedwell, D.M. Aminoglycoside antibiotics restore CFTR function by overcoming premature stop mutations. Nat Med 2, 467–469 (1996).

4. Welch, E.M. et al. PTC124 targets genetic disorders caused by nonsense mutations. Nature 447, 87–91 (2007).

5. Albers, S. et al. Engineered tRNAs suppress nonsense mutations in cells and in vivo. Nature 618, 842–848 (2023).

6. Porter, J.J., Heil, C.S. & Lueck, J.D. Therapeutic promise of engineered nonsense suppressor tRNAs. Wiley Interdiscip Rev RNA 12, e1641 (2021).

7. Nishikura, K. A-to-I editing of coding and non-coding RNAs by ADARs. Nat Rev Mol Cell Biol 17, 83–96 (2016).

8. Reautschnig, P. et al. CLUSTER guide RNAs enable precise and efficient RNA editing with endogenous ADAR enzymes in vivo. Nat Biotechnol 40, 759–768 (2022).

9. Katrekar, D. et al. Efficient in vitro and in vivo RNA editing via recruitment of endogenous ADARs using circular guide RNAs. Nat Biotechnol 40, 938–945 (2022).

10. Yi, Z. et al. Engineered circular ADAR-recruiting RNAs increase the efficiency and fidelity of RNA editing in vitro and in vivo. Nat Biotechnol 40, 946–955 (2022).

11. Eggington, J.M., Greene, T. & Bass, B.L. Predicting sites of ADAR editing in double-stranded RNA. Nat Commun 2, 319 (2011).

12. Borchardt, E.K., Martinez, N.M. & Gilbert, W.V. Regulation and Function of RNA Pseudouridylation in Human Cells. Annu Rev Genet 54, 309–336 (2020).

13. Kufel, J. & Grzechnik, P. Small Nucleolar RNAs Tell a Different Tale. Trends Genet 35, 104–117 (2019).

14. Karijolich, J. & Yu, Y.T. Converting nonsense codons into sense codons by targeted pseudouridylation. Nature 474, 395–398 (2011).

15. Song, J. et al. CRISPR-free, programmable RNA pseudouridylation to suppress premature termination codons. Mol Cell 83, 139–155 e139 (2023).

16. Adachi, H. et al. Targeted pseudouridylation: An approach for suppressing nonsense mutations in disease genes. Mol Cell 83, 637–651 e639 (2023).

17. Cong, L. et al. Multiplex genome engineering using CRISPR/Cas systems. Science 339, 819–823 (2013).

18. Mali, P. et al. RNA-guided human genome engineering via Cas9. Science 339, 823–826 (2013).

19. Gadgil, A. & Raczynska, K.D. U7 snRNA: A tool for gene therapy. J Gene Med 23, e3321 (2021).

20. Rogalska, M.E. et al. Therapeutic activity of modified U1 core spliceosomal particles. Nat Commun 7, 11168 (2016).

21. Raitskin, O., Cho, D.S., Sperling, J., Nishikura, K. & Sperling, R. RNA editing activity is associated with splicing factors in lnRNP particles: The nuclear pre-mRNA processing machinery. Proc Natl Acad Sci U S A 98, 6571–6576 (2001).

22. Flanigan, K.M. et al. Nonsense mutation-associated Becker muscular dystrophy: interplay between exon definition and splicing regulatory elements within the DMD gene. Hum Mutat 32, 299–308 (2011).

23. Love, M.I., Huber, W. & Anders, S. Moderated estimation of fold change and dispersion for RNA-seq data with DESeq2. Genome Biol 15, 550 (2014).

24. Zhou, Y. et al. Metascape provides a biologist-oriented resource for the analysis of systems-level datasets. Nat Commun 10, 1523 (2019).

25. Vaquero-Garcia, J. et al. RNA splicing analysis using heterogeneous and large RNA-seq datasets. Nat Commun 14, 1230 (2023).

26. Patel, S.B. & Bellini, M. The assembly of a spliceosomal small nuclear ribonucleoprotein particle. Nucleic Acids Res 36, 6482–6493 (2008).

27. Ngo, L.H. et al. Nuclear export of circular RNA. Nature 627, 212–220 (2024).

28. Hsiao, Y.E. et al. RNA editing in nascent RNA affects pre-mRNA splicing. Genome Res 28, 812–823 (2018).

29. Cao, Y. et al. RNA-based translation activators for targeted gene upregulation. Nat Commun 14, 6827 (2023).

30. Choi, J. et al. 2’-O-methylation in mRNA disrupts tRNA decoding during translation elongation. Nat Struct Mol Biol 25, 208–216 (2018).

31. Elliott, B.A. et al. Modification of messenger RNA by 2’-O-methylation regulates gene expression in vivo. Nat Commun 10, 3401 (2019).

32. Arango, D. et al. Direct epitranscriptomic regulation of mammalian translation initiation through N4-acetylcytidine. Mol Cell 82, 2797–2814 e2711 (2022).

33. Thalalla Gamage, S., et al. Antisense pairing and SNORD13 structure guide RNA cytidine acetylation. RNA 28, 1582–1596 (2022).

