## Supplemental Information for "Small nuclear RNAs enhance protein-free RNA-programmable base conversion on mammalian coding transcripts"

*Supplementary Information*

Aaron A. Smargon[1,2,3], Deepak Pant[1,2,3], Sofia Glynne[1,2,3], Trent A. Gomberg[1,2,3], Gene W. Yeo[1,2,3,*]

[1] Department of Cellular and Molecular Medicine, University of California San Diego, 9500 Gilman Drive, La Jolla, CA 92093 USA

[2] Stem Cell Program, University of California San Diego, Sanford Consortium for Regenerative Medicine, 2880 Torrey Pines Scenic Drive, La Jolla, CA 92037 USA

[3] Institute for Genomic Medicine, University of California San Diego, 9500 Gilman Drive, La Jolla, CA 92093 USA

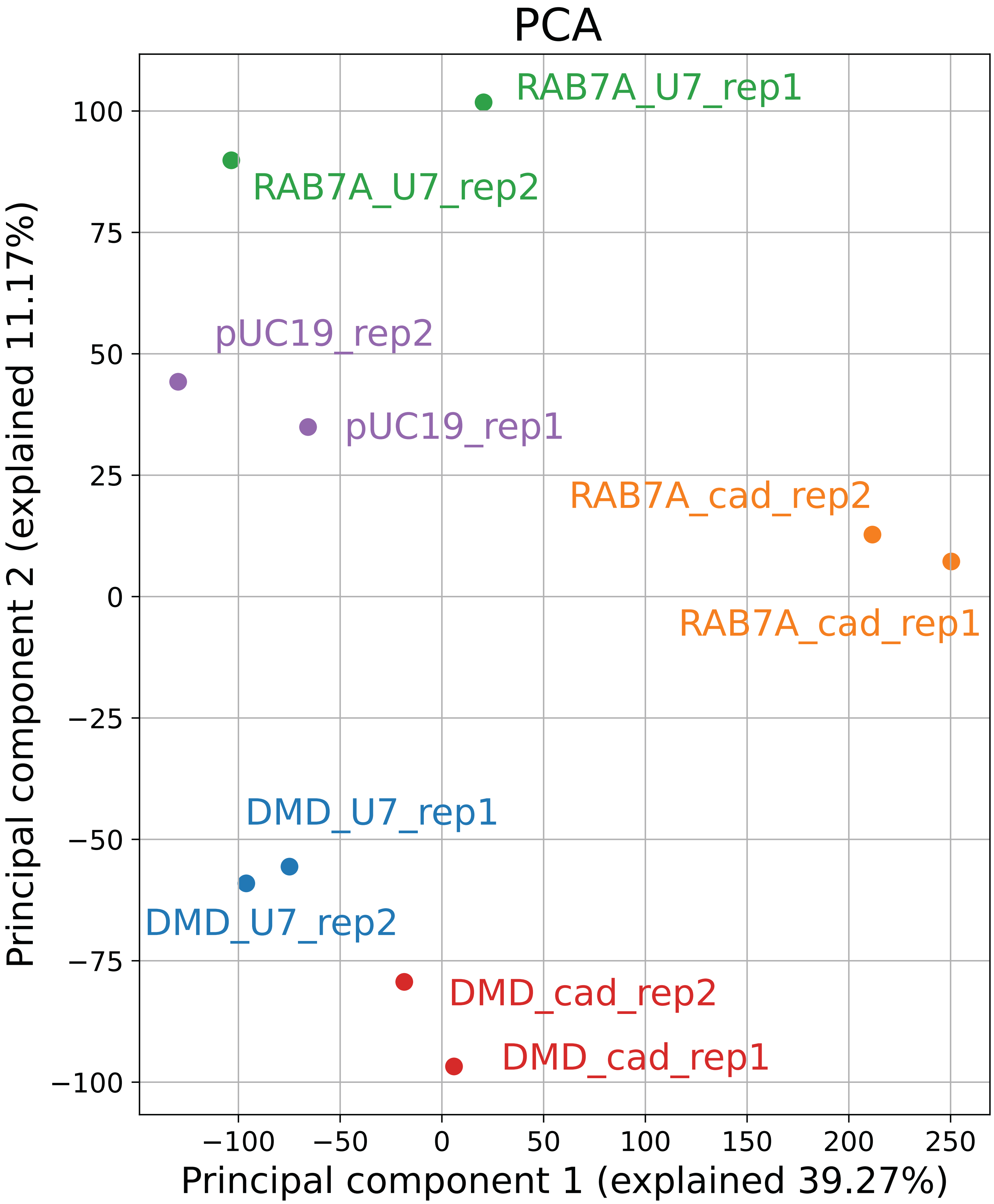


### Supplementary Figure 1

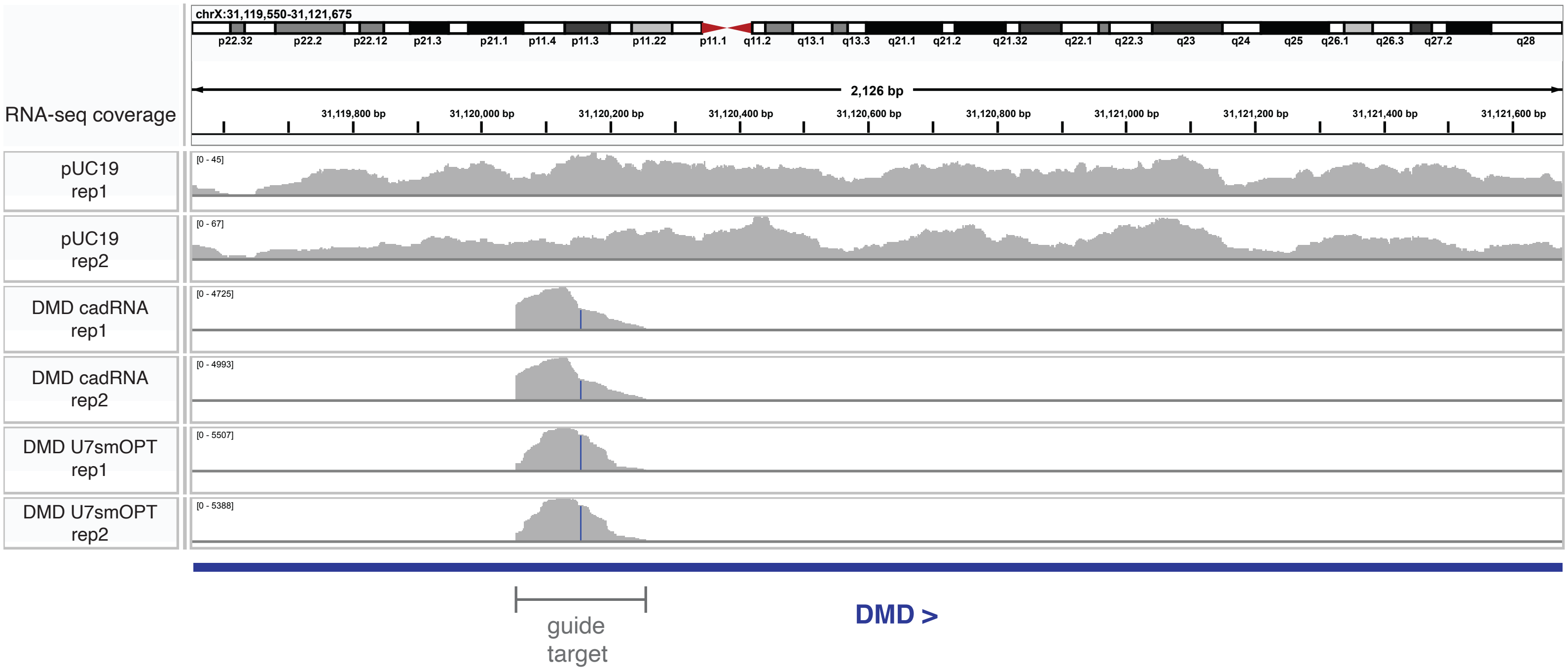


### Supplementary Figure 2

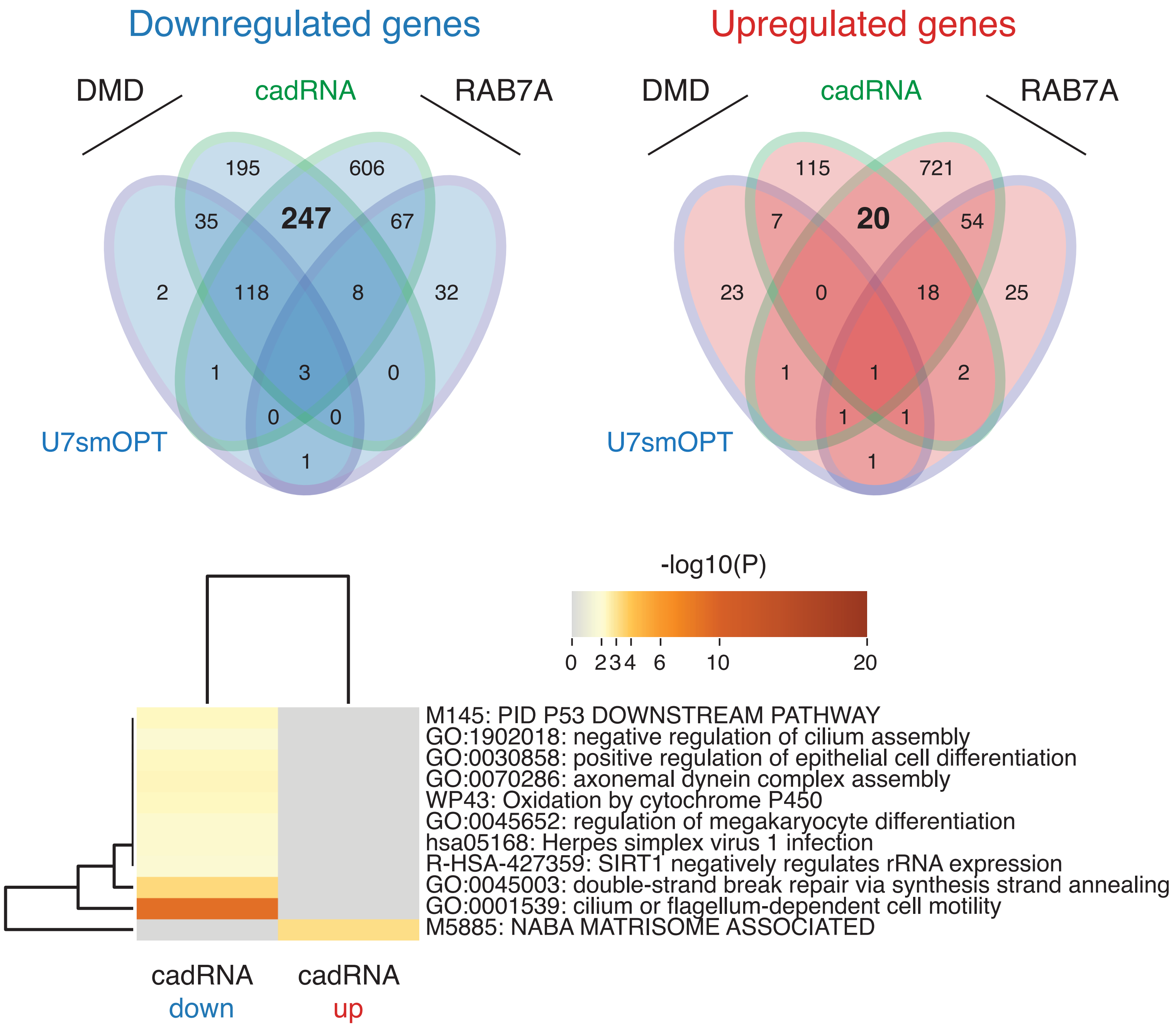


### Supplementary Figure 3

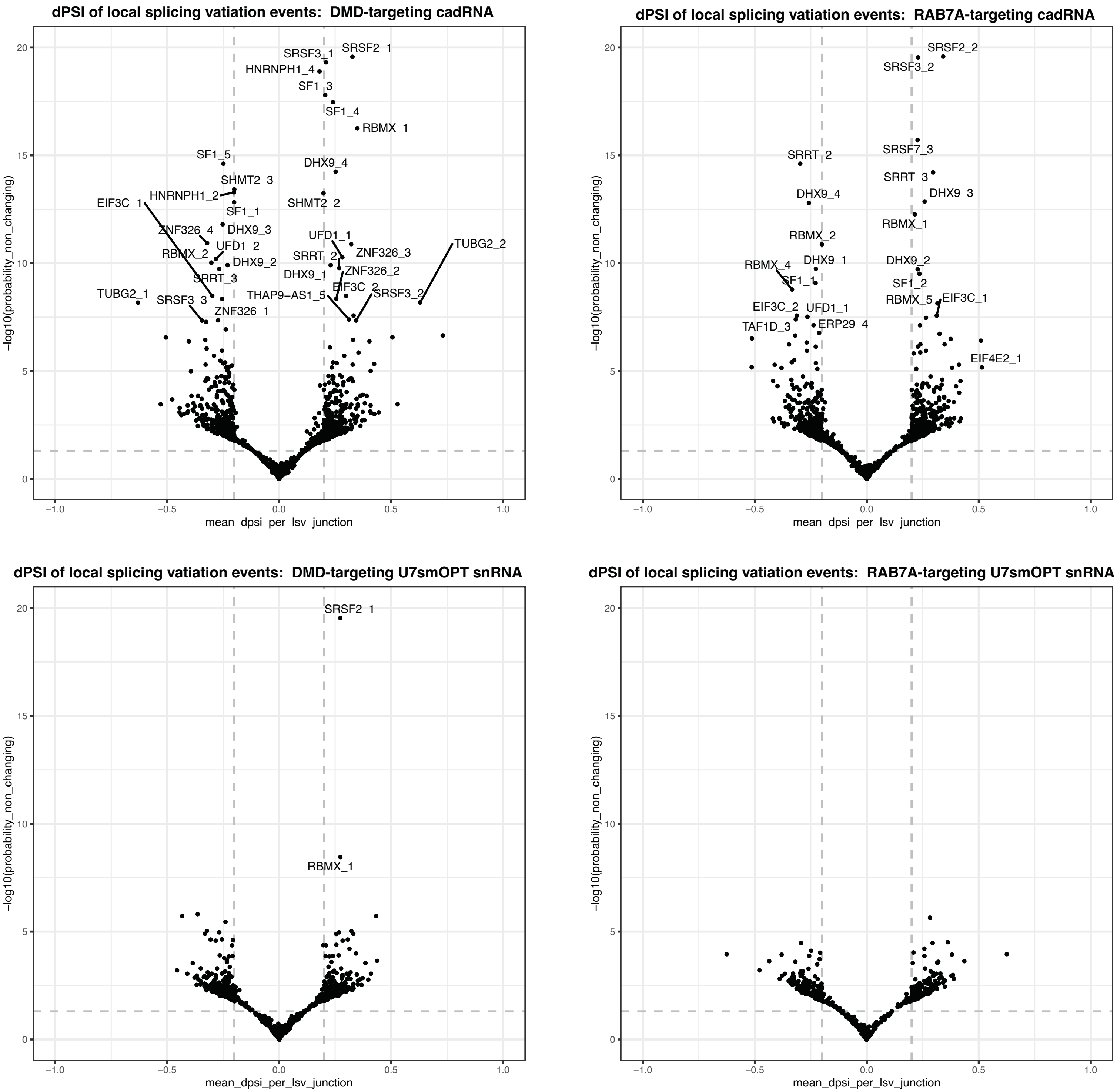


### Supplementary Figure 4

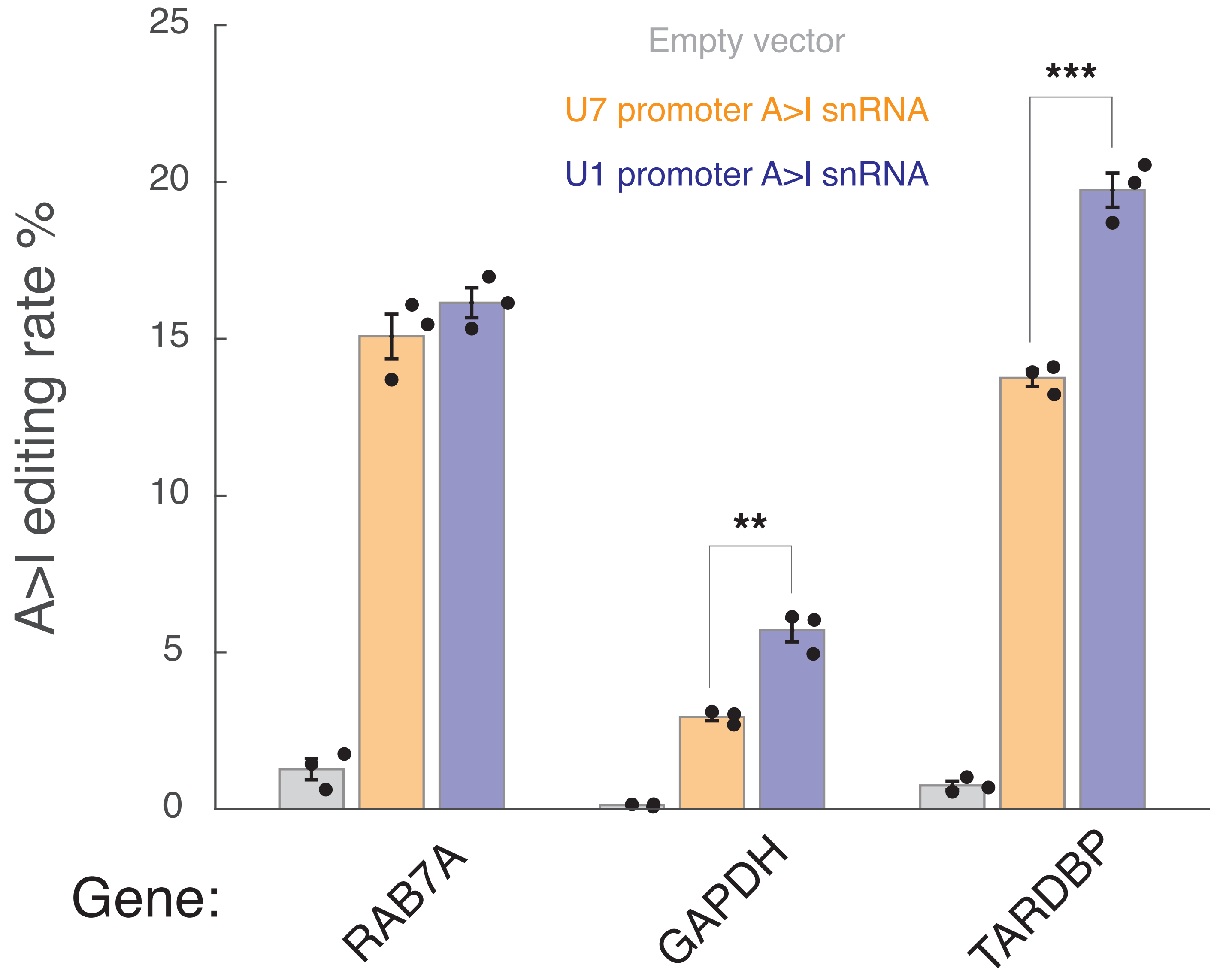


### Supplementary Figure 5

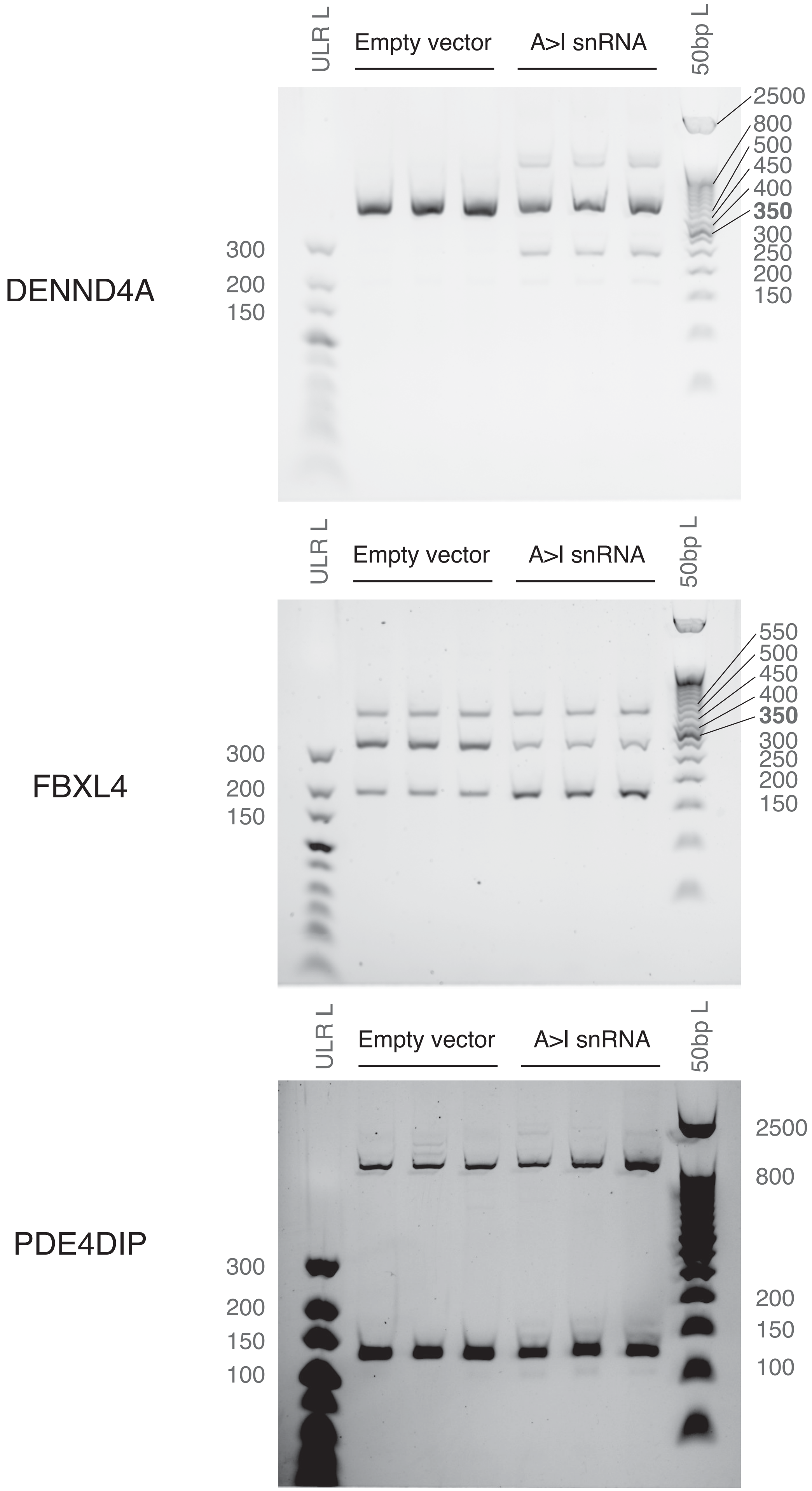


### Supplementary Figure 6

#### SUPPLEMENTARY FIGURE LEGENDS

**Supplementary Figure 1: Principal Component Analysis (PCA) plot of RNA-guided A>I base converter RNA sequencing data.** PCA plot of RNA sequencing samples analyzed in Figure 2, with each of two replicates colored by sample (pUC19, RAB7A-targeting cadRNA, RAB7A-targeting U7smOPT snRNA, DMD-targeting cadRNA, and DMD-targeting U7smOPT snRNA).

**Supplementary Figure 2: Alignment of RNA-guided A>I base converter RNA sequencing reads to DMD gene.** Alignment showing RNA sequencing read pile-up across DMD-targeting cadRNA and U7smOPT snRNA replicates at the guide target on DMD.

**Supplementary Figure 3: Pathway analysis of significantly perturbed genes by RNA-guided A>I base converters.** 4-way Venn diagrams of the number of significantly downregulated and upregulated genes across conditions from Figure 2 (top). Enriched pathway heatmap of downregulated and upregulated genes conserved across both cadRNA guides (bottom).

**Supplementary Figure 4: Differential splicing analysis of RNA-guided A>I base converter RNA sequencing data.** Scatterplots of local splicing variations (LSVs) against empty control (pUC19) of cadRNA backbone vs. U7smOPT snRNA backbone for RAB7A- and DMD-targeting guides. Cutoffs for significance are $p$-value $< 0.05$ and various dPSI (differential Percent Spliced In) values used in Figure 2.

**Supplementary Figure 5: Editing performance of A>I snRNAs with U7 vs. U1 snRNA promoter.** Editing percent performance by transfection in HEK293T cells of A>I snRNAs targeting three different genes and driven by either U7 or U1 snRNA promoter. Overperformance significance vs. U7 promoter: **, ***: $p$ < 1e-2, 1e-3 (one-way ANOVA). Error bars reflect standard error of mean.

**Supplementary Figure 6: Splicing gels for A>I snRNA-targeted pre-mRNA.** Full RT-PCR gels for all replicates of experiments in Figure 3.

**SUPPLEMENTARY TABLES**

<u>**Supplementary Table 1: Plasmid sequences.**</u>

| **Name** | **Sequence** |
|---:|---|
| **U1<br>promoter-<br>terminator-<br>U1_snRNA<br>backbone** | taacacaggctaaggaccagcttctttgggagagaacagacgcaggggcgggagggaaaaagggagaggca<br>gacgtcacttccccttggcggctctggcagcagattggtcggttgagtggcagaaaggcagacggggactggg<br>caaggcactgtcggtgacatcacggacagggcgacttctatgtagatgaggcagcgcagaggctgctgcttcgc<br>cacttgctgcttcaccacgaaggagttcccgtgccctgggagcgggttcaggaccgctgatcggaagtgagaat<br>cccagctgtgtgtcagggctggaaagggctcgggagtgcgcggggcaagtgaccgtgtgtgtaaagagtgag<br>gcgtatgaggctgtgtcggggcagaggcccaagatctcA**<guide>**gcaggggagataccatgatcacgaa<br>ggtggttttcccagggcgaggcttatccattgcactccggatgtgctgacccctgcgatttccccaaatgtgggaa<br>actcgactgcataatttgtggtagtgggggactgcgttcgcgctttcccctgactttctggagtttcaaaagtagact<br>gtacgctaa |
| **U7<br>promoter-<br>terminator-<br>U7smOPT_<br>snRNA<br>backbone** | ggcttaacaacaacgaaggggctgtgactggctgctttctcaaccaatcagcaccgaactcatttgcatgggctga<br>gaacaaatgttcgcgaactctagaaatgaatgacttaagtaagttccttagaatattatttttcctactgaaagttacca<br>catgcgtcgttgtttatacagtaataggaacaagaaaaaagtcacctaagctcaccctcatcaattgtggagttccttt<br>atatcccatcttctctccaaacacatacgcagC**<guide>**agaatttTtGGagtaggctttctggctttttaccgg<br>aaagcccctcttatgatgtttgttgccaatgatagattgttttcactgtgcaaaaattatgggtagttttggtggtcttgat<br>gcagttgtaagcttggggtatgaaggtttgggccacgcctggg |
| **U1<br>promoter-<br>terminator-<br>U7smOPT_<br>snRNA<br>backbone** | taacacaggctaaggaccagcttctttgggagagaacagacgcaggggcgggagggaaaaagggagaggca<br>gacgtcacttccccttggcggctctggcagcagattggtcggttgagtggcagaaaggcagacggggactggg<br>caaggcactgtcggtgacatcacggacagggcgacttctatgtagatgaggcagcgcagaggctgctgcttcgc<br>cacttgctgcttcaccacgaaggagttcccgtgccctgggagcgggttcaggaccgctgatcggaagtgagaat<br>cccagctgtgtgtcagggctggaaagggctcgggagtgcgcggggcaagtgaccgtgtgtgtaaagagtgag<br>gcgtatgaggctgtgtcggggcagaggcccaagatctcA**<guide>**agaatttTtGGagtaggctttctggc<br>tttttaccggaaagcccctactttctggagtttcaaaagtagactgtacgctaa |
| **U6<br>promoter-<br>terminator-<br>cadRNA<br>backbone** | gagggcctatttcccatgattccttcatatttgcatatacgatacaaggctgttagagagataattagaattaatttgac<br>tgtaaacacaaagatattagtacaaaatacgtgacgtagaaagtaataatttcttgggtagtttgcagttttaaaattat<br>gttttaaaatggactatcatatgcttaccgtaacttgaaagtatttcgatttcttggctttatatatcttgtggaaaggac<br>Gaaacaccgccatcagtcgccggtcccaagcccggataaaatgggagggggcgggaaaccgcctaaccatg<br>ccgactgatggcagaAAAAAAAAA**<guide>**AAAAAAAAAActgccatcagtcggcgtgg<br>actgtagaacactgccaatgccggtcccaagcccggataaaagtggagggtacagtccacgctttttt |
| **U6<br>promoter-<br>terminator-<br>for-<br>snoRNAs<br>backbone** | gagggcctatttcccatgattccttcatatttgcatatacgatacaaggctgttagagagataattagaattaatttgac<br>tgtaaacacaaagatattagtacaaaatacgtgacgtagaaagtaataatttcttgggtagtttgcagttttaaaattat<br>gttttaaaatggactatcatatgcttaccgtaacttgaaagtatttcgatttcttggctttatatatcttgtggaaaggac<br>G**<snoRNA>**tttttt |
| **U7<br>promoter-<br>terminator-<br>for-U>Ψ-<br>snRNAs<br>backbone** | ggcttaacaacaacgaaggggctgtgactggctgctttctcaaccaatcagcaccgaactcatttgcatgggctga<br>gaacaaatgttcgcgaactctagaaatgaatgacttaagtaagttccttagaatattatttttcctactgaaagttacca<br>catgcgtcgttgtttatacagtaataggaacaagaaaaaagtcacctaagctcaccctcatcaattgtggagttccttt<br>atatcccatcttctctccaaacacatacgcagC**<U>ΨsnRNA>**cttatgatgtttgttgccaatgatagattgt<br>tttcactgtgcaaaaattatgggtagttttggtggtcttgatgcagttgtaagcttggggtatgaaggtttgggccacg<br>cctggg |

| | |
|---|---|
| **CFTR PTC dual luciferase reporter** | GACATTGATTATTGACTAGTTATTAATAGTAATCAATTACGGGGTCATT<br>AGTTCATAGCCCATATATGGAGTTCCGCGTTACATAACTTACGGTAAAT<br>GGCCCGCCTGGCTGACCGCCCAACGACCCCCGCCCATTGACGTCAAT<br>AATGACGTATGTTCCCATAGTAACGCCAATAGGGACTTTCCATTGACG<br>TCAATGGGTGGAGTATTTACGGTAAACTGCCCACTTGGCAGTACATCA<br>AGTGTATCATATGCCAAGTACGCCCCCTATTGACGTCAATGACGGTAA<br>ATGGCCCGCCTGGCATTATGCCCAGTACATGACCTTATGGGACTTTCC<br>TACTTGGCAGTACATCTACGTATTAGTCATCGCTATTACCATGGTGATG<br>CGGTTTTGGCAGTACATCAATGGGCGTGGATAGCGGTTTGACTCACG<br>GGGATTTCCAAGTCTCCACCCCATTGACGTCAATGGGAGTTTGTTTTG<br>GCACCAAAATCAACGGGACTTTCCAAAATGTCGTAACAACTCCGCCC<br>CATTGACGCAAATGGGCGGTAGGCGTGTACGGTGGGAGGTCTATATA<br>AGCAGAGCTCTCTGGCTAACTAGAGAACCCACTGCTTACTGGCTTAT<br>CGAAATTAATACGACTCACTATAGGGAGACCCAAGCTGGCTAGCGTT<br>TAAACGGGCCCTCTAGACTCGAGCGGCCGCCACTGTGCTGGATATCT<br>GCAGGCCACCatggcttccaaggtgtacgaccccgagcaacgcaaacgcatgatcactgggcctcagt<br>ggtgggctcgctgcaagcaaatgaacgtgctggactccttcatcaactactatgattccgagaagcacgccgaga<br>acgccgtgatttttctgcatggtaacgctgcctccagctacctgtggaggcacgtcgtgcctcacatcgagcccgt<br>ggctagatgcatcatccctgatctgatcggaatgggtaagtccggcaagagcgggaatggctcatatcgcctcct<br>ggatcactacaagtacctcaccgcttggttcgagctgctgaaccttccaaagaaaatcatctttgtgggccacgact<br>gggggcttgtctggcctttcactactcctacgagcaccaagacaagatcaaggccatcgtccatgctgagagtg<br>tcgtggacgtgatcgagtcctgggacgagtggcctgacatcgaggaggatatcgccctgatcaagagcgaaga<br>gggcgagaaaatggtgcttgagaataacttcttcgtcgagaccatgctcccaagcaagatcatgcggaaactgga<br>gcctgaggagttcgctgcctacctggagccattcaaggagaagggcgaggttagacggcctaccctctcctggc<br>ctcgcgagatccctctcgttaagggaggcaagcccgacgtcgtccagattgtccgcaactacaacgcctaccttc<br>gggccagcgacgatctgcctaagatgttcatcgagtccgaccctgggttcttttccaacgctattgtcgagggagc<br>taagaagttccctaacaccgagttcgtgaaggtgaagggcctccacttcagccaggaggacgctccagatgaaa<br>tgggtaagtacatcaagagcttcgtggagcgcgtgctgaagaacgagcagaattctgcttgcaagaactggttca<br>gtagcttaagccactttgtgatccaccttaacagccacggcttccctcccgaggtggaggagcaggccgccggc<br>accctgcccatgagctgcgcccaggagagcggcatggatagacaccctgctgcttgcgccagcgccaggatca<br>acgtcAGCGGAAGTGAGACACCGGGTACGAGTGAGTCAGCTACTCCAG<br>AAAGT**gattcaataactttgcaacagtgAaggaaagcctttggagtgata**GGTTCTgccgatgct<br>aagaacattaagaagggccctgctcccttctaccctctggaggatggcaccgctggcgagcagctgcacaaggc<br>catgaagaggtatgccctggtgcctggcaccattgccttcaccgatgcccacattgaggtggacatcacctatgcc<br>gagtacttcgagatgtctgtgcgcctggccgaggccatgaagaggtacggcctgaacaccaaccaccgcatcgt<br>ggtgtgctctgagaactctctgcagttcttcatgccagtgctgggcgccctgttcatcggagtggccgtggcccct<br>gctaacgacatttacaacgagcgcgagctgctgaacagcatgggcatttctcagcctaccgtggtgttcgtgtcta<br>agaagggcctgcagaagatcctgaacgtgcagaagaagctgcctatcatccagaagatcatcatcatggactcta<br>agaccgactaccagggcttccagagcatgtacacattcgtgacatctcatctgcctcctggcttcaacgagtacga<br>cttcgtgccagagtctttcgacagggacaaaaccattgccctgatcatgaacagctctgggtctaccggcctgcct<br>aagggcgtggccctgcctcatcgcaccgcctgtgtgcgcttctctcacgcccgcgaccctattttcggcaaccag<br>atcatccccgacaccgctattctgagcgtggtgccattccaccacggcttcggcatgttcaccaccctgggctacc<br>tgatttgcggctttcgggtggtgctgatgtaccgcttcgaggaggagctgttcctgcgcagcctgcaagactacaa<br>aattcagtctgccctgctggtgccaaccctgttcagcttcttcgctaagagcaccctgatcgacaagtacgacctgt<br>ctaacctgcacgagattgcctctggcggcgccccactgtctaaggaggtgggcgaagccgtggccaagcgcttt<br>catctgccaggcatccgccagggctacggcctgaccgagacaaccagcgccattctgattaccccagagggcg |

| | |
|---|---|
| | acgacaagcctggcgccgtgggcaaggtggtgccattcttcgaggccaaggtggtggacctggacaccggca<br>agaccctgggagtgaaccagcgcggcgagctgtgtgtgcgcggccctatgattatgtccggctacgtgaataac<br>cctgaggccacaaacgccctgatcgacaaggacggctggctgcactctggcgacattgcctactgggacgagg<br>acgagcacttcttcatcgtggaccgcctgaagtctctgatcaagtacaagggctaccaggtggccccagccgag<br>ctggagtctatcctgctgcagcaccctaacattttcgacgccggagtggccggcctgcccgacgacgatgccgg<br>cgagctgcctgccgccgtcgtcgtgctggaacacggcaagaccatgaccgagaaggagatcgtggactatgtg<br>gccagccaggtgacaaccgccaagaagctgcgcggcggagtggtgttcgtggacgaggtgcccaagggcct<br>gaccggcaagctggacgcccgcaagatccgcgagatcctgatcaaggctaagaaaggcggcaagatcgccgt<br>gtaaGATCCGAGCTCGGTACCAAGCTTAAGTTTAAACCGCTGATCAGCC<br>TCGACTGTGCCTTCTAGTTGCCAGCCATCTGTTGTTTGCCCCTCCCCC<br>GTGCCTTCCTTGACCCTGGAAGGTGCCACTCCCACTGTCCTTTCCTAA<br>TAAAATGAGGAAATTGCATCGCATTGTCTGAGTAGGTGTCATTCTATT<br>CTGGGGGGTGGGGTGGGGCAGGACAGCAAGGGGGAGGATTGGGAA<br>GACAATAGCAGGCATGCTGGGGATGCGGTGGGCTCTATGG |

**Supplementary Table 2: Guide and snoRNA sequences.**

| Name | Sequence |
|---:|---|
| **RAB7A A>I guide** | AGACAGTTGTCCCCCTGGAGAGATGAAATCGATGTTGGCTCTTA<br>ATGGAAAGATAAAAGGCGTACATTCAAGATGTGTCTACTGTACA<br>GAATACTGCCGCCAGCTGCTAATCCCAATTCTGAGTATGTGTCTG<br>CAATCCAAACACCCATCAACCCTCCACCTTTGTCGCCTGCATTAC<br>AGGAGAATAACACATAATCCAA |
| **DAXX A>I guide** | GCTCCTGTAACCTGATGCCCACATCTCGGAAGGCATCCTGAGCC<br>ATGAGCTGGAGCTGCTGTCGGGGGAGGCCAAGGCTGTGTCGGG<br>CAGCTGCCTTCTCCACAGCCCGAAGCACATCCCCATAGTCAGGG<br>AAGGTATCAGGCCCTGGCTTGTTGATGAGCCGCTCAATGCGCCT<br>GTTAACCTCTGGGTAGCGGGTGCCA |
| **GAPDH A>I guide** | GGCCATCCACAGTCTTCTGGGTGGCAGTCTACGCATGGACTGTG<br>GTCTACTGTCCTTCCACGATACGTTCGTTGTCATGGATGACCTTG<br>GCCAGGGGTGCCAAGCACAACGTGGTGCAGGAGGCAAACGTG<br>ATGATCTTGAGGCACAAGTCATACTTCTCATGCAAGACACCCATG<br>ACGAACATGGGGGCATCAGCAGAG |
| **TARDBP A>I guide** | TGTGTTTCATATTCCGTAAAACGAACAATCGGAAACCCCTTTGA<br>ATGTGGTGTCTTAAGATCTTTCAACTCCTGCACCATAAGAACTTC<br>TCCAAAGGTACCAAAATTGAGTTTCAGGTCCTGTTCCCAAGTGT<br>TCCATGGGAGAGGGTACACTATTAAATCGGTACAGTTCTGGACT<br>GCTCTTGTCACTTTCACTGCTGA |
| **ALDOA A>I guide** | GTCCGTCCTTCTTGTACTGGGCACAGCGCTCAGACAGCCCATCC<br>AACCCTTGGGTGGTAGTCTCGCCATTTGTCCCTGCCAGGGGGAC<br>CACGCCCTTGTCCACCTTGATGCCCACAACACCGCCCTTGGATT<br>TGATAACTTGGGGGAAGGGACGCCCATCATCCGCCTTCTGGTAG<br>AGTGTCTCATGGAAGAGGATGACAC |
| **SMAD4 A>I guide** | TTGTAGTCCACCATCCTGATAAGGTTAACCCGCCCAACGGTAAA<br>AGAGGAGAGTCTAAAGGTTGTGCCAGTGCAATCGGCATGGTATG<br>AAGTACTTCGTCCAGGAGGACCAGGGCCCGGTGTAAGACGGTT<br>TCAATCCAGCAAGCACATTCTTTGATGCTCTGAGAAGGGTAATC<br>CGGTCCCCAGCCTTTCACAAAACTC |
| **FANCC A>I guide** | GAGTCTGGGCTGAGGGACCTGGCTCTGCTAAATGTAAAATAGAT<br>ACTTCGTGATTGTCCCAAGATGACATCAGCTCATTCTCACAGCCC<br>AGCGAGGGCACCTACTCGTGTAATGCGTGGCCACAGCAGTTCAC<br>CTGTCCTGTGGGGGAGGCGAGCCTGATCCCAGTGGCCGGGCAC<br>CCACACGGCCTGCGTGCCTTCTAG |
| **DMD A>I guide** | aaagctaattacacttgatgtcagcccactctccaaaagctaattacacttgatgtcagaggtaacagatttg<br>caaaattataggtcacacggtggtatcCattgaatgaatgatttaaaaatcaaaaagaaataaaatggcatg<br>aaagagtaaagctttttcctaccagtccttagcttttcctcttgagcttttctcct |
| **MDN1 A>I guide** | acaataaaatttttgtagttgtccaaaagggagcacctgggtaagcaatgtgaccttctgaccacagttaag<br>tctcactttggactcttctttctgttcCatgggtggtcagaggctgtcaccaactcaaaccactgtctgaggg<br>catcgctgagtgtctcaggaagtgcgtttacatctcgaagaatgatatagtatgg |

| | |
|---|---|
| **UBR4 A>I guide** | acaactcatggctcctaggtatgtacaggccctttgatggcttgggttacagacaacctcatagctggtgcaccacacacacgagataaaacaggaagccCaaaaaccccaagccacaccaagaaaaatgagagagggggagggcggggtaacaatgcagcatcccgcggagggaacttaatgcacaaggagggagaacaga |
| **DENND4A A>I guide** | gatgaggtactcccactaatatttgcggggcgcccatagggagtactctgaataatttcacaaccctgtttcaatctttctttccagtcatataaaacccCaaaaataaatttttaaaaataaatacacaaagcactttaaatatgaaaagtaatattctcaggcatcatcaactgatccttaaatgtagtagcatggtaa |
| **FBXL4 A>I guide** | aggaagactggagactcctatccagcttcaaagcttcttgaaaatccaagcgaatgaagcaaatggagaaggcaagggaaccaggccaggaagaaaaatcCaaaagcagagcaaaaatctcaaaaaccagaggtaaacagtaccactctcttcgaggaatcaaggtagatctcaaaaatgcaaggaaataatccaaaaagt |
| **PDE4DIP A>I guide** | gagcctgaagtcttcgattgtcttccctgaggactctgttctcattgcagagctgaggtatggactccaggccctgactgtagaagttagaagtggatccCgtcccagaaagtagacatttgggtaacagaaactttcaaaagtttcccattcgttatctctcaaagatactgccccaacccagcagaaaaccccaaggac |
| **IDUA H/ACA snoRNA** | GTGCACATcttcgccGACCTGCTTTCTTCTATGTGAGTAGTGTgactgcATGTGCTATACAAATAATTGAAGGctttggtcGCAGTATAACTATAAATAGTAATGCTGCgagttgcCCTTCAGACAAAA |
| **CFTR H/ACA snoRNA** | GTGCACATCTTTCCTTGACCTGCTTTCTTCTATGTGAGTAGTGTCCTGTTGCATGTGCTATACAAATAATTGAAGGCTTTCCTTGCAGTATAACTATAAATAGTAATGCTGCCTGTTCCCTTCAGACAAAA |
| **ACTB-U1226 H/ACA snoRNA** | GTGCACATAACGCAACGACCTGCTTTCTTCTATGTGAGTAGTGTCAGTCATAATGTGCTATACAAATAATTGAAGGACGCAACGCAGTATAACTATAAATAGTAATGCTGCAGTCAACCTTCAGACAAAA |
| **EEF2-U2881 H/ACA snoRNA** | GTGCACATTAAGTCCCGACCTGCTTTCTTCTATGTGAGTAGTGTCCTAAGAGATGTGCTATACAAATAATTGAAGGAAGTCCCGCAGTATAACTATAAATAGTAATGCTGCCTAAGACCTTCAGACAAAA |
| **RPS6-U514 H/ACA snoRNA** | GTGCACATGGCTTTCTGACCTGCTTTCTTCTATGTGAGTAGTGTTCAACATAATGTGCTATACAAATAATTGAAGGCCTTTCTGCAGTATAACTATAAATAGTAATGCTGCCAACATCCTTCAGACAAAA |

**Supplementary Table 3: Primers for PCR, qPCR, Sanger sequencing, and NGS.**

| Name | Sequence |
|---:|---|
| **RAB7A A>I FOR** | CCTCCCTCCTTGAAGGCTACC |
| **RAB7A A>I REV** | AAGCTCCGCTAACCTAAGAATACC |
| **GAPDH A>I FOR** | GATGCTGGCGCTGAGTACGT |
| **GAPDH A>I REV** | CACCACTGACACGTTGGCAGT |
| **TARDBP A>I FOR** | CCATCGGAAGACGATGGGACG |
| **TARDBP A>I REV** | CCTGAATGGCTTGGGGATGAAG |
| **FANCC A>I FOR** | CCTGCACAACAGCTGATCAGGC |
| **FANCC A>I REV** | TTCTTTAATGGTTCATGACCAAATTCTTGG |
| **ALDOA A>I FOR** | GACAGCTGACGACCGCGTGAACC |
| **ALDOA A>I REV** | CCCCAATCTTCAGCACACAACGCCA |
| **DAXX A>I FOR** | CTGACCGGCCGTGTCATAGAGCAGC |
| **DAXX A>I REV** | GTGAGGTGGCAGCCAAAGTTGTAGATGA |
| **SMAD4 A>I FOR** | GCGTGCACCTGGAGATGCTG |
| **SMAD4 A>I REV** | ACAGGTGAAGAATTAATAAGAATGTGTTTCTCCT |
| **MDN1 A>I FOR** | cagaggaagaccaagacccc |
| **MDN1 A>I REV** | AAAACCTCAGCCCAGGCAAA |
| **UBR4 A>I FOR** | GAGTACATCCGCCACAACGA |
| **UBR4 A>I REV** | agtatgaagctggccggga |
| **DMD A>I FOR** | GCGAGTAGTTCCACACAGGT |
| **DMD A>I REV** | TCAGGAACACCCCAAAACCA |
| **NEAT1 qPCR FOR** | GCTGGACCTTTCATGTAACGGG |
| **NEAT1 qPCR REV** | TGAACTCTGCCGGTACAGGGAA |
| **GAPDH qPCR FOR** | GTCTCCTCTGACTTCAACAGCG |
| **GAPDH qPCR REV** | ACCACCCTGTTGCTGTAGCCAA |
| **RAB7A guide qPCR FOR** | AGTTGTCCCCCTGGAGAGATG |
| **RAB7A guide qPCR REV** | CCTGTAATGCAGGCGACAAA |
| **GAPDH guide qPCR FOR** | CGCATGGACTGTGGTCTACT |
| **GAPDH guide qPCR REV** | CCATGTTCGTCATGGGTGTC |
| **TARDBP guide qPCR FOR** | GAGTTTCAGGTCCTGTTCCCA |
| **TARDBP guide qPCR REV** | AAGTGACAAGAGCAGTCCAGAA |
| **DENND4A A>I FOR** | GCCAAAATGCCCAGTTGCTT |
| **DENND4A A>I REV** | TACTCATTGCTTTTGTAGTGGGC |
| **FBXL4 A>I FOR** | TTCCTTCAGTTCCTTTTTCTGCTT |
| **FBXL4 A>I REV** | AGTGTCAGCATGGAAACTACTCA |
| **PDE4DIP A>I FOR** | GTAAGGGTCCTCCCACCTCT |
| **PDE4DIP A>I REV** | GCTGTGAGTTTGGGTTGGTG |
| **DENND4A RT-PCR FOR** | TTACCCCAACTGGATTGTCAG |
| **DENND4A RT-PCR REV** | TACAGTGTTCGTCTTTGCCAC |
| **FBXL4 RT-PCR FOR** | TTGTGCCGCATCTAGAGAGTC |
| **FBXL4 RT-PCR REV** | CCAGGTACCCTGAACTTTGC |
| **PDE4DIP RT-PCR FOR** | GAACACCGGCTGACCTCTAC |

| | |
|---|---|
| **PDE4DIP RT-PCR REV** | CTTCGATTGTCTTCCCTGAGGAC |
| **ACTB-U1226-TAG-F** | ACACTCTTTCCCTACACGACGCTCTTCCGATCTNNNNNt ccatcgtccaccgcaaatgct |
| **ACTB-U1226-TAG-R** | TGGAGTTCAGACGTGTGCTCTTCCGATCTNNNNNNgcc aatctcatcttgttttctgcgca |
| **EEF2-U2881-TAG-F** | ACACTCTTTCCCTACACGACGCTCTTCCGATCTNNNNNc agagtccggaggcagcag |
| **EEF2-U2881-TAG-R** | TGGAGTTCAGACGTGTGCTCTTCCGATCTNNNNNNaaa gtgttgggtgtcccatccc |
| **RPS6-U514-TAA-F** | ACACTCTTTCCCTACACGACGCTCTTCCGATCTNNNNNc gcaaacttttcaatctctcTAAa |
| **RPS6-U514-TAA-R** | TGGAGTTCAGACGTGTGCTCTTCCGATCTNNNNNNagg acacgtggagtaacaag |
| **Universal NGS Primer 1** | AATGATACGGCGACCACCGAGATCTACACTCTTTCCCT ACACGACGCTCTTCCGATCT |
| **Index NGS Primer 2-1** | CAAGCAGAAGACGGCATACGAGATCATACCACGTGAC TGGAGTTCAGACGTGTGCTCTTCCGATC |
| **Index NGS Primer 2-2** | CAAGCAGAAGACGGCATACGAGATGAAGTTGGGTGA CTGGAGTTCAGACGTGTGCTCTTCCGATC |
| **Index NGS Primer 2-3** | CAAGCAGAAGACGGCATACGAGATATGACGTCGTGAC TGGAGTTCAGACGTGTGCTCTTCCGATC |
| **Index NGS Primer 2-4** | CAAGCAGAAGACGGCATACGAGATTTGGACGTGTGA CTGGAGTTCAGACGTGTGCTCTTCCGATC |
| **Index NGS Primer 2-5** | CAAGCAGAAGACGGCATACGAGATAGTGGATCGTGA CTGGAGTTCAGACGTGTGCTCTTCCGATC |
| **Index NGS Primer 2-6** | CAAGCAGAAGACGGCATACGAGATGATAGGCTGTGA CTGGAGTTCAGACGTGTGCTCTTCCGATC |
| **Index NGS Primer 2-7** | CAAGCAGAAGACGGCATACGAGATTGGTAGCTGTGA CTGGAGTTCAGACGTGTGCTCTTCCGATC |
| **Index NGS Primer 2-8** | CAAGCAGAAGACGGCATACGAGATCGCAATCTGTGAC TGGAGTTCAGACGTGTGCTCTTCCGATC |
| **Index NGS Primer 2-9** | CAAGCAGAAGACGGCATACGAGATGATGTGTGGTGA CTGGAGTTCAGACGTGTGCTCTTCCGATC |
| **Index NGS Primer 2-10** | CAAGCAGAAGACGGCATACGAGATGATTGCTCGTGAC TGGAGTTCAGACGTGTGCTCTTCCGATC |
| **Index NGS Primer 2-11** | CAAGCAGAAGACGGCATACGAGATCGCTCTATGTGAC TGGAGTTCAGACGTGTGCTCTTCCGATC |
| **Index NGS Primer 2-12** | CAAGCAGAAGACGGCATACGAGATTATCGGTCGTGAC TGGAGTTCAGACGTGTGCTCTTCCGATC |
| **Index NGS Primer 2-13** | CAAGCAGAAGACGGCATACGAGATAACGTCTGGTGA CTGGAGTTCAGACGTGTGCTCTTCCGATC |
| **Index NGS Primer 2-14** | CAAGCAGAAGACGGCATACGAGATACGTTCAGGTGA CTGGAGTTCAGACGTGTGCTCTTCCGATC |
| **Index NGS Primer 2-15** | CAAGCAGAAGACGGCATACGAGATCAGTCCAAGTGA CTGGAGTTCAGACGTGTGCTCTTCCGATC |

| | |
|---|---|
| **Index NGS Primer 2-16** | CAAGCAGAAGACGGCATACGAGATTTGCAGACGTGACTGGAGTTCAGACGTGTGCTCTTCCGATC |
| **Index NGS Primer 2-17** | CAAGCAGAAGACGGCATACGAGATCAATGTGGGTGACTGGAGTTCAGACGTGTGCTCTTCCGATC |
| **Index NGS Primer 2-18** | CAAGCAGAAGACGGCATACGAGATACTCCATCGTGACTGGAGTTCAGACGTGTGCTCTTCCGATC |
| **Index NGS Primer 2-19** | CAAGCAGAAGACGGCATACGAGATGTTGACCTGTGACTGGAGTTCAGACGTGTGCTCTTCCGATC |
| **Index NGS Primer 2-20** | CAAGCAGAAGACGGCATACGAGATCGTGTGTAGTGACTGGAGTTCAGACGTGTGCTCTTCCGATC |
| **Index NGS Primer 2-21** | CAAGCAGAAGACGGCATACGAGATACGACTTGGTGACTGGAGTTCAGACGTGTGCTCTTCCGATC |
| **Index NGS Primer 2-22** | CAAGCAGAAGACGGCATACGAGATCACTAGCTGTGACTGGAGTTCAGACGTGTGCTCTTCCGATC |
| **Index NGS Primer 2-23** | CAAGCAGAAGACGGCATACGAGATACTAGGAGGTGACTGGAGTTCAGACGTGTGCTCTTCCGATC |
| **Index NGS Primer 2-24** | CAAGCAGAAGACGGCATACGAGATGTAGGAGTGTGACTGGAGTTCAGACGTGTGCTCTTCCGATC |
| **Index NGS Primer 2-25** | CAAGCAGAAGACGGCATACGAGATCCTGATTGGTGACTGGAGTTCAGACGTGTGCTCTTCCGATC |
| **Index NGS Primer 2-26** | CAAGCAGAAGACGGCATACGAGATATGCACGAGTGACTGGAGTTCAGACGTGTGCTCTTCCGATC |
| **Index NGS Primer 2-27** | CAAGCAGAAGACGGCATACGAGATCGACGTTAGTGACTGGAGTTCAGACGTGTGCTCTTCCGATC |
| **Index NGS Primer 2-28** | CAAGCAGAAGACGGCATACGAGATTACGCCTTGTGACTGGAGTTCAGACGTGTGCTCTTCCGATC |
| **Index NGS Primer 2-29** | CAAGCAGAAGACGGCATACGAGATCCGTAAGAGTGACTGGAGTTCAGACGTGTGCTCTTCCGATC |
| **Index NGS Primer 2-30** | CAAGCAGAAGACGGCATACGAGATATCACACGGTGACTGGAGTTCAGACGTGTGCTCTTCCGATC |
| **Index NGS Primer 2-31** | CAAGCAGAAGACGGCATACGAGATCACCTGTTGTGACTGGAGTTCAGACGTGTGCTCTTCCGATC |
| **Index NGS Primer 2-32** | CAAGCAGAAGACGGCATACGAGATCTTCGACTGTGACTGGAGTTCAGACGTGTGCTCTTCCGATC |
| **Index NGS Primer 2-33** | CAAGCAGAAGACGGCATACGAGATTGCTTCCAGTGACTGGAGTTCAGACGTGTGCTCTTCCGATC |
| **Index NGS Primer 2-34** | CAAGCAGAAGACGGCATACGAGATAGAACGAGGTGACTGGAGTTCAGACGTGTGCTCTTCCGATC |
| **Index NGS Primer 2-35** | CAAGCAGAAGACGGCATACGAGATGTTCTCGTGTGACTGGAGTTCAGACGTGTGCTCTTCCGATC |
| **Index NGS Primer 2-36** | CAAGCAGAAGACGGCATACGAGATTCAGGCTTGTGACTGGAGTTCAGACGTGTGCTCTTCCGATC |
